# SLBP-independent control of maternal histone mRNA

**DOI:** 10.64898/2026.01.06.697898

**Authors:** Joana Pereirinha, Martin Brehm, Shamitha Govind, Anke Busch, Nadezda Podvalnaya, Ann-Sophie Seistrup, Kamila Delaney, Florian A Steiner, Julian Konig, Sebastian Falk, René F. Ketting

**Affiliations:** Biology of Non-coding RNA group, Institute of Molecular Biology, Ackermannweg 4, 55128 Mainz, Germany; International PhD Programme on Gene Regulation, Epigenetics & Genome Stability, Mainz, Germany; Max Perutz Labs, Vienna Biocenter Campus (VBC), Dr.-Bohr-Gasse 9, 1030 Vienna, Austria; University of Vienna, Max Perutz Labs, Department of Structural and Computational Biology, Campus Vienna Biocenter 5, 1030 Vienna, Austria; Vienna Biocenter PhD Program, a Doctoral School of the University of Vienna and the Medical University of Vienna, 1030 Vienna, Austria; Bioinformatics Core Facility, Institute of Molecular Biology, Ackermannweg 4, 55128 Mainz, Germany; CECAD Cluster of Excellence, Institute for Genetics, CECAD Research Center, Joseph-Stelzmann-Str. 26, 50931 Cologne, Germany; Department of Molecular and Cellular Biology, Section of Biology, Faculty of Sciences, University of Geneva, Geneva 1211, Switzerland; RNA Modifications & Regulation group, Institute of Molecular Biology, Ackermannweg 4, 55128 Mainz, Germany; Theodor Boveri Institute, Biocenter, University of Würzburg, Am Hubland, 97074 Würzburg, Germany; Institute of Developmental Biology and Neurobiology, Johannes Gutenberg University, Mainz, Germany

## Abstract

Replication-dependent (RD) histones are crucial for packaging newly replicated DNA into chromatin, ensuring genome stability. In metazoans, the mRNA of RD histones is uniquely regulated through a conserved 3′ stem–loop bound by stem–loop binding protein (SLBP). This allows cell cycle-coupled regulation of these important transcripts. However, oocytes must stabilise histone mRNAs independently of the cell cycle to ensure maternal loading to support the first embryonic divisions. Using *Caenorhabditis elegans* as a model system, we discovered an SLBP-independent mechanism that ensures RD histone transcript stability during oogenesis. This is mediated by the protein complex PETISCO, bound to the effector protein TOST-1, which directly binds the histone stem–loop region and maintains maternal histone mRNA levels during oogenesis and early embryogenesis. Loss of this mechanism disrupts histone homeostasis, leading to premature genome activation, mitotic defects, and embryonic lethality. Interestingly, the same complex, PETISCO, acts in piRNA biogenesis when bound to the effector PID-1, revealing an intriguing co-option of this histone mRNA homeostasis mechanism by the piRNA pathway. Our findings reveal a unique SLBP-independent mechanism of histone mRNA regulation, that served as a basis for the evolution of a novel piRNA biogenesis mechanism.

## Introduction

The very early stages of embryonic development rely on the availability of maternally provided materials, as the zygotic genome itself is not yet active. Only after processes known as the maternal-to-zygotic transition (MZT)^1^ and zygotic genome activation (ZGA)^2^ can the embryo produce products from its own genome, making it independent of the maternally provided material.

Among the most critical maternal components are the core histone proteins: their quantity must be tightly balanced with DNA to ensure chromatin integrity and appropriate transcriptional timing^3–7^. Perturbations in histone supply disrupt this balance: excess histones delay ZGA and extend nuclear divisions, while histone depletion accelerates transcription onset, lengthens cell cycles, and triggers checkpoint activation^7–12^.

Replication-dependent (RD) histones are encoded by a distinct class of eukaryotic genes whose mRNAs are uniquely non-polyadenylated in metazoans, terminating instead in a conserved 3′ stem-loop structure essential for post-transcriptional regulation^13,14^. Stem-loop binding protein (SLBP) binds this element and orchestrates histone mRNA processing, nuclear export, translation, and degradation^15–23^. In proliferating somatic cells, histone mRNA levels are tightly coupled to the cell cycle, peaking during S phase, followed by rapid degrading afterwards^14^. However, in the germline, notably during oogenesis, this paradigm is bypassed: histone mRNAs are transcribed and stabilised in a cell cycle–independent manner to ensure their maternal deposition into the embryo^24,25^.

To meet the high demand for histone supply during early embryogenesis, organisms have evolved diverse strategies to stockpile histone mRNAs and/or proteins^14^. In *Drosophila*, histone mRNAs are transcribed in nurse cells at the end of oogenesis and stored via SLBP-dependent mechanisms; in addition, histone proteins are stored ^8,24–27^. In *Xenopus*, where genome size and the number of pre-ZGA divisions are greater, both histone proteins and mRNAs accumulate early in oogenesis; an oocyte-specific SLBP variant binds histone mRNAs to repress translation until fertilisation^17,20,28,29^. In zebrafish, an oocyte-specific SLBP is again required for RD histone mRNA storage^30^. Additionally, an eIF4E1 paralogue has been found to selectively bind non/lowly-polyadenylated transcripts, including RD histones, likely contributing to their stabilisation in the early embryo^31^. In *Caenorhabditis elegans*, RD histone mRNAs share conserved features with those of other metazoans, including the essential 3′ stem-loop structure^32,33^. The sole SLBP homolog, termed CDL-1, specifically binds this hairpin and is essential for post-transcriptional regulation of histone gene expression, including facilitating 3′ end processing and controlling translation^34,35^. However, not much is known about how these nematodes regulate their maternal histone mRNA pool.

PETISCO, a protein complex comprising PID-3, ERH-2, TOFU-6, and IFE-3, was initially characterised for its role in piRNA biogenesis in *C. elegans*^36–38^. PETISCO binds and stabilises piRNA precursors, for which it requires the binding of effector protein PID-1 via ERH-2^37–39^. However, instead of binding PID-1, PETISCO can also bind an alternative effector, TOST-1, in a mutually exclusive manner. This reprograms PETISCO towards a distinct, piRNA-independent function. TOST-1 does not affect piRNAs but is required maternally to support embryogenesis^37,38^. Interestingly, TOST-1 is more deeply conserved than PID-1, suggesting the TOST-1-related function of PETISCO represents PETISCO’s ancestral function^37^.

Here, we uncover the critical function for the PETISCO:TOST-1 complex during embryogenesis: it regulates histone mRNA accumulation during oogenesis in the adult *C. elegans* germline. We demonstrate that PETISCO:TOST-1 specifically binds and stabilises RD histone mRNAs via their 3′ stem-loop region, enabling maternal histone mRNA accumulation and proper embryonic development. Notably, this mechanism operates independently of SLBP and reveals PETISCO:TOST-1 as the first known SLBP-independent regulator of RD-histone mRNAs. These findings also provide a unique example of how an existing gene-regulatory module, histone mRNA stabilization, can be adopted into a new role, piRNA biogenesis.

## Results

### TOST-1 and PETISCO are conserved factors required maternally for embryonic viability

TOST-1 is essential for *C. elegans* embryonic development, as demonstrated by the maternal-effect lethality (Mel) phenotype observed in *tost-1* loss-of-function mutants (Figure 1a,b). In the *tost-1(xf194)* null allele, which harbours a 445 bp deletion disrupting the first exon and eliminating ERH-2 binding residues, 100% of embryos from homozygous mutant mothers failed to develop (Figure 1a). A hypomorphic allele, *tost-1(xf196 ts)*, leading to a C-terminal truncation while preserving the ERH-2 interaction domain, also caused complete embryonic lethality at 25°C but showed intermediate viability at 20°C and was fully viable at 15°C (Figure 1b). This temperature sensitivity suggests that residual TOST-1 function becomes critical under stress conditions or that the truncated protein may be destabilised at elevated temperatures. Embryos from *tost-1(xf196 ts)* mutant mothers grown at 25°C exhibited mitotic abnormalities, including chromatin bridges and micronuclei formation. This indicates defective chromosome segregation during early stages of cell division, resulting in severe defects in later stages (Figures 1c, Extended Data Fig 1a). Notably, *pid-1* mutations partially rescued the *tost-1(xf196 ts)* temperature-sensitive phenotype at 25°C but had no effect on the null *tost-1(xf194)* allele (Extended Data Fig 1b,c). This genetic interaction suggests that in hypomorphic conditions, an increase in available PID-1-free PETISCO may compensate for reduced TOST-1 function.

**Figure 1.**
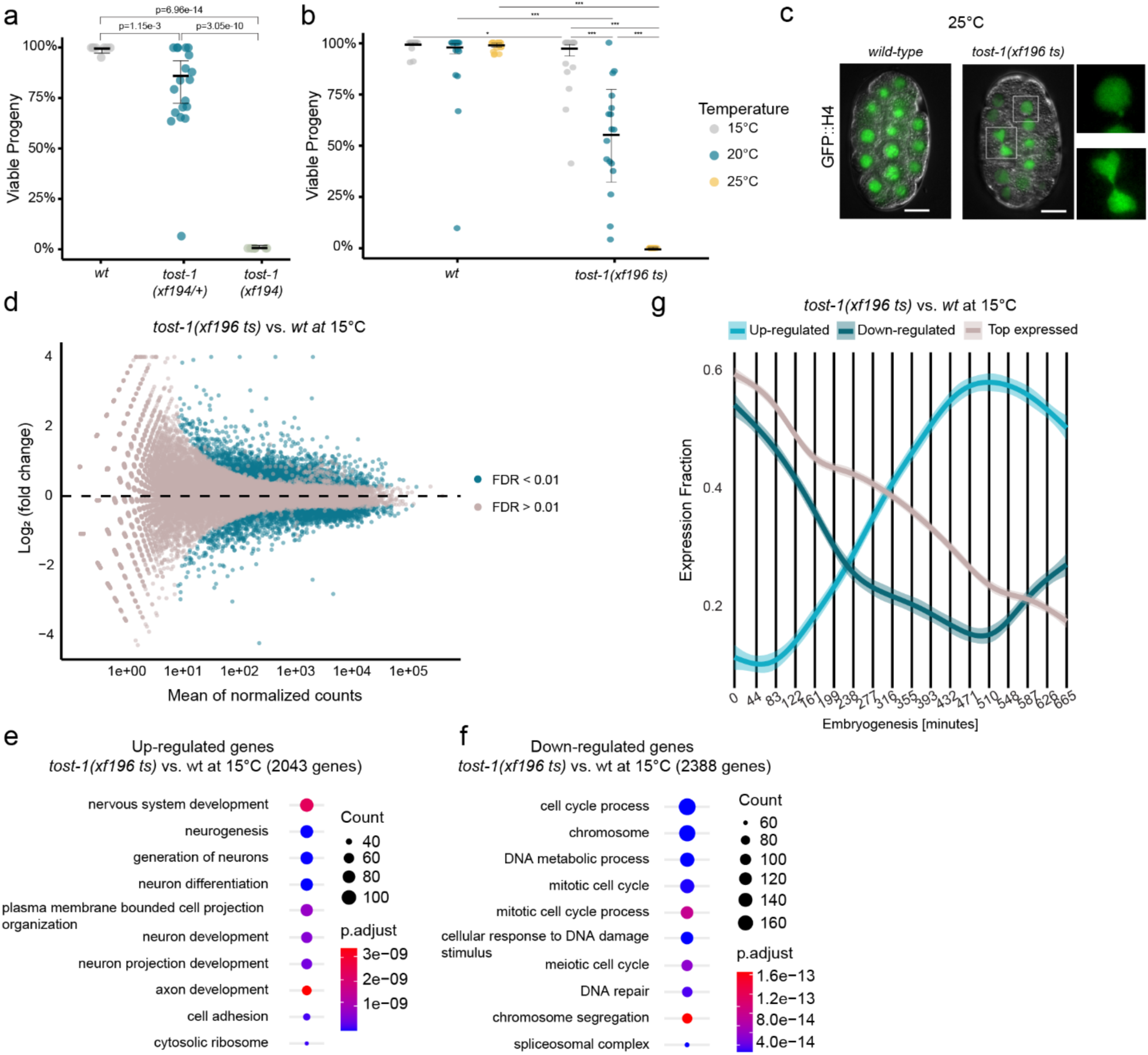
TOST-1 is essential for embryonic viability and proper gene expression. **a,** Embryonic viability of *wild-type* (n=8), *tost-1(xf194/+)* heterozygous (n=19), and *tost-1(xf194)* homozygous mutant (n=9) mothers. Statistical analysis using maximum a posteriori estimation within a Bayesian Generalised Linear Mixed-Effects Model with Tukey method for multiple comparisons. Error bars represent 95% confidence intervals of the estimated viability. **b,** Temperature-sensitive embryonic lethality in *tost-1(xf196 ts)* mutants. Embryonic viability at 15°C, 20°C, and 25°C for *wild-type* (n=20, 18, 19 respectively) and *tost-1(xf196 ts)* (n=19, 17, 19 respectively) mothers. Statistical analysis using maximum a posteriori estimation within a Bayesian Generalised Linear Mixed-Effects Model with specific pairwise comparisons. Error bars represent 95% confidence intervals of the estimated viability. P values were corrected using the multivariate-t method within each temperature. Asterisks indicate significant differences: *P<0.05, **P<0.01, ***P<0.001. Exact P values are indicated in Extended Data Table 1. **c,** Representative images of *C. elegans* embryos showing mitotic defects in *tost-1(xf196 ts)* mutants. Overlay of fluorescence and brightfield images of transgene *xfSi254*[GFP::H4] in *wild-type* (left) or *tost-1(xf196 ts)* (right) background grown at 25°C. Insets showing mitotic defects. Scale bar, 10 μm. **d,** Differential expression MA plot of RNA-seq analysis comparing gene expression in *tost-1(xf196 ts)* versus *wild-type* embryos at 15°C. Blue dots indicate significantly upregulated and downregulated genes (FDR <0.01). Adjusted P values calculated using Benjamini-Hochberg method. n = 3 biological replicates. **e,f,** Gene Ontology (GO) term enrichment analysis of upregulated (**e**) and downregulated (**f**) genes from panel d. Top 10 enriched GO terms shown for biological processes, molecular functions, and cellular components. Adjusted P values calculated using Benjamini-Hochberg method (FDR <0.05). **g,** Temporal expression patterns during wild-type embryogenesis. Expression fraction (y-axis) plotted against embryogenesis time in minutes (x-axis) for three gene sets: top 25% expressed genes (in all conditions) from panel d, upregulated genes in *tost-1* mutants, and downregulated genes in *tost-1* mutants. Shaded areas represent 95% confidence intervals. Data derived from publicly available wild-type embryogenesis dataset^40^.

To test whether PETISCO’s role in embryonic development extends beyond *C. elegans*, we examined the complex in the closely related nematode *C. briggsae*, which diverged 80-110 million years ago, an evolutionary distance comparable to that between human and mouse. All PETISCO subunits have clear homologs, and immunoprecipitation confirmed that the complex assembles similarly in both species (Extended Data Fig 1d). Yeast two-hybrid analysis revealed largely conserved protein-protein interactions, with only the PID-3/TOFU-6 interaction failing to be detected (Extended Data Fig 1e). Functionally, RNAi depletion of *C. briggsae* PID-3 and TOFU-6 produced the same Mel phenotype observed in *C. elegans*, demonstrating that PETISCO’s essential developmental role is evolutionarily conserved in nematodes (Extended Data Fig 1f).

### TOST-1 affects the timing of gene expression in embryos

To understand the molecular basis of TOST-1’s essential role, we performed RNA-sequencing on early embryos (isolated from gravid mothers) from *tost-1(xf196 ts)* mutants and *wild-type* controls at permissive (15°C) and restrictive (25°C) temperatures. Already at 15°C, where embryonic lethality is minimal, *tost-1* mutants showed extensive transcriptional dysregulation: 2,043 genes were upregulated and 2,388 downregulated compared to *wild-type* embryos (Figure 1d). At the restrictive temperature (25°C), dysregulation was also severe, with 2,499 upregulated and 2,457 downregulated genes (Extended Data Fig 1g). However, since the 25°C data may include secondary effects of embryonic arrest (Extended Data Fig 1h,i), we focused our analysis on the 15°C dataset to identify primary transcriptional changes. Gene ontology analysis revealed a clear pattern: upregulated genes were enriched for neurogenesis functions, while downregulated genes were associated with cell cycle progression (Figure 1e,f). Furthermore, comparison with published maternal and zygotic expression profiles^40^ revealed that the upregulated genes are normally expressed later in embryogenesis, while the downregulated genes correspond to maternal and early zygotic transcripts (Figure 1g). This pattern resembles the effect of premature ZGA.

To further probe whether TOST-1 regulates the timing of ZGA, we monitored GFP::H4 expression using the *xfSi268* reporter transgene, in which the *his-67* promoter drives a GFP coding sequence (lacking introns) fused to *his-67* and its endogenous 3′ UTR. This transgene is not expressed in the germline but only starts to be expressed during embryogenesis, making it a useful readout for ZGA timing (Extended Data Fig 2a).

RNAi-mediated depletion of *tost-1* or the PETISCO component *tofu-6* resulted in earlier GFP::H4 expression compared to controls, consistent with the idea of premature ZGA in the absence of PETISCO:TOST-1 (Extended Data Fig 2b). In zebrafish, ZGA is promoted by the depletion of maternal histones^6^. Hence, we aimed to test if direct histone knockdown may cause a comparable effect on our reporter. Indeed, RNAi targeting *his-65* (encoding histone H2A) produced a similar phenotype of earlier transgene activation (Extended Data Fig 2b). Using the quantification of nuclear GFP intensity to fit a linear model, and establishing the mean intensity of nuclear GFP at timepoint 4 as the detection limit, we found that *tofu-6* and *his-65* RNAi embryos had significantly higher expression at the onset of activation (intercept of the linear model).This is also evident at timepoint 10, where the control remained below the detection limit (Extended Data Fig 2c,d). For *tost-1* RNAi, GFP nuclear intensity was above the wild-type signal but did not reach statistical significance, likely due to the low number of replicates (n=2). Importantly, the regression slopes did not differ significantly between treatments, indicating that once transcription begins, the rate of GFP::H4 expression increase is similar across all conditions. This suggests that the RNAi treatments affect the *timing* of transgene activation rather than the overall transcriptional rate, consistent with premature ZGA (Extended Data Fig 2c). This precocious transcriptional activation was accompanied by delayed cell divisions: both *his-65* and *tofu-6* RNAi embryos exhibited reduced cell numbers at later stages compared to control RNAi (Extended Data Fig 2d). These phenotypes are consistent with previous observations that histone depletion activates checkpoint responses while paradoxically accelerating ZGA^5,7^. Since histone availability modulates the onset of transcription by altering the histone-to-DNA ratio, these findings raise the possibility that TOST-1 may regulate ZGA timing through histone mRNA regulation.

### TOST-1 regulates RD histone mRNA stability in the germline

Given that PETISCO depletion phenocopies histone depletion, we next examined whether histone transcript levels were altered in *tost-1* mutants. We quantified *his-65* and *his-66* mRNA (encoding histones H2A and H2B, respectively) in young adults and embryos at different temperatures using RT–qPCR. Due to sequence similarity among histone genes, the primers used to amplify *his-65* and *his-66* also recognize other, but not all, histone gene copies, providing an overall indication of histone gene expression. Notably, in young adults, both *tost-1* alleles consistently showed significant reduction in *H2A* transcript abundance at all temperatures and similar, but weaker effects on *H2B* (Extended Data Fig 3a), indicating that PETISCO:TOST-1 is needed for maintaining histone mRNA levels in this stage. In embryos, we observed a big variation in the expression of both transcript pools, even in the *wild-type* control (indicated by the error bars, representing the fold change variation relative to *wild-type*). This is likely due to the usage of pooled embryos with different embryonic stages. However, we generally observe a reduction of histone transcripts, with *H2B* expression being significantly reduced at 20°C in both *tost-1* mutants (Extended Data Fig 3b).

To visualise histone mRNA expression with spatial resolution, we performed single-molecule FISH (smFISH) in adult animals using probes for *his-16* and *his-60* (encoding H2A and H4, respectively) that showed specific signal in their respective fluorescence channels (Extended Data Fig 3c). Again, due to the high sequence similarity among histone gene families, these probes likely cross-react with multiple histone genes, providing a representative measure of H2A and H4 histone gene expression.

To visualize histone mRNA distribution in the germline, we initially used GFP::WAGO-3^41^ to mark gonadal tissue in *wild-type* and *tost-1(xf196 ts)* mutant animals. This revealed a strong reduction of *H4* smFISH signal in *tost-1(xf196 ts)* mutants (Extended Data Fig 3d-f). Because quantification yielded equivalent results whether the region of interest (ROI), the germline, was defined using GFP::WAGO-3 or *H4* smFISH signal, we proceeded with the latter. Similar effects were detected using probes targeting *H2A* (Figure 2a,b). In *wild-type* animals, the staining intensities appeared to drop after the mitotic region, consistent with histone requirement during mitosis, and to increase around the gonadal turn, where germ cells are in the diplotene stage (Figure 2a and Extended Data Fig 3g). This suggests that at this stage of meiosis, non-S-phase histone gene transcription takes place. Notably, by calculating the fold change of the signal in the meiotic *vs* the mitotic region, we observed a significant reduction of signal in the meiotic region in *tost-1* mutants for *H2A*. Also with the *H4* probes a decrease in expression, especially in the oocytes, was evident (Figure 2a,c). The H4 decrease did not reach statistical significance, likely because of the accumulation of punctate signals around the gonadal turn. Stronger reductions in histone transcript abundance were observed in *tost-1(xf194)* and *pid-3(tm2417)* mutants (Figure 2d-i). Interestingly, in the various mutants, but especially in *tost-1(xf196 ts)*, we detected *H2A/H4* mRNA granules around the gonadal bend, the region where in wild-type animals overall intensities increased. We hypothesise that these result from S-phase independent transcription starting during the diplotene phase of meiosis (see Discussion).

**Figure 2.**
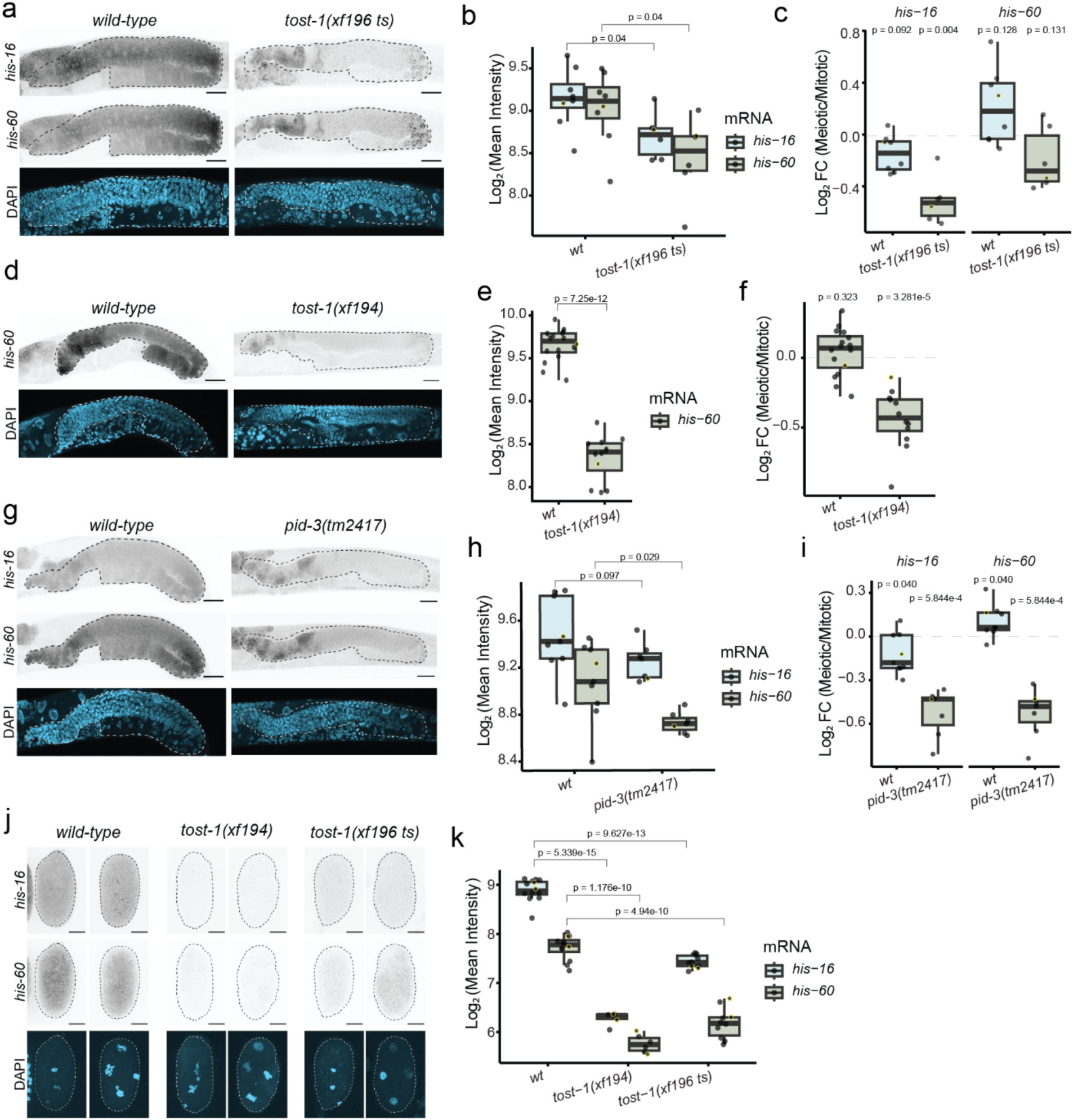
TOST-1 regulates histone mRNA levels in the germline and early embryos. **a,** Representative smFISH images of *wild-type* and *tost-1(xf196 ts)* young adult gonads. Top: inverted fluorescence image of *his-16* probes; middle: inverted fluorescence image of *his-60* probes; bottom: DAPI fluorescence. Scale bars, 20 μm. **b,** Quantification of histone smFISH signal (log2 mean intensity) in gonads of *wild-type* (n=8) and *tost-1(xf196 ts)* (n=6) animals. P values calculated using Welch’s t-test with Holm correction for multiple comparisons. **c,** Log2 fold change (FC) of histone smFISH signal in meiotic versus mitotic regions from samples in panel b. P values calculated using paired t-test with Holm correction. **d,** Representative smFISH images of *wild-type* (n=16) and *tost-1(xf194)* (n=12) gonads using *his-60* probes. **e,f**, Quantification of *his-60* smFISH signal (e) and meiotic/mitotic fold change (f) as in panels b,c. **e,** P value calculated using Welch’s t-test. **f,** P values calculated using paired t-test with Holm correction. **g,** Representative smFISH images of *wild-type* (n=9) and *pid-3 (tm2417)* (n=7) mutant gonads using *his-16* and *his-60* probes (inverted fluorescence). **h,i,** Quantification of histone smFISH signal (h) and meiotic/mitotic fold change (i) as in panels b,c. **h,** P values calculated using Welch’s t-test with Holm correction. **i,** P values calculated using paired t-test with Holm correction. **j,** Representative smFISH images of 2-cell and 4-cell stage embryos from *wild-type* (n=12), *tost-1(xf194)* (n=6), and *tost-1(xf196 ts)* (n=10) using *his-16* and *his-60* probes (inverted fluorescence). Scale bars, 10 μm. **k,** Quantification of *his-16* and *his-60* smFISH signal in embryos. Statistical analysis using Welch’s t-test with Holm correction. Yellow data points indicate samples corresponding to representative images shown.

Since *C. elegans* zygotic transcription begins after the 4-cell stage^42,43^, we also examined histone mRNA levels in embryos up to this stage, in order to specifically assess maternal histone mRNA levels in embryos. Both *tost-1* mutant alleles exhibited significant reductions in *H2A* and *H4* mRNA signal (Figure 2j,k). Together, these results demonstrate that PETISCO:TOST-1 is required to stabilise histone transcripts during oogenesis, ensuring maternal deposition for the support of early embryogenesis.

### Genetic interaction between a histone cluster and *tost-1*

To genetically test whether histone mRNA depletion underlies the *tost-1* developmental phenotype, we examined genetic interactions between histone gene dosage and TOST-1 function. As described above, the hypomorphic *tost-1(xf196 ts)* allele produces viable, fertile animals at permissive temperatures (15°C and 20°C). We generated a large chromosomal deletion removing 13 clustered histone genes from chromosome II (*his-9* through *his-16*, plus *his-25*, *his-26*, and *his-42* through *his-44*), *ugeDf12* (Figure 3a), and combined it with *tost-1(xf196 ts)*. While *ugeDf12* animals alone are fully fertile at all temperatures, the *tost-1(xf196 ts); ugeDf12* double mutants exhibited complete maternal-effect lethality, even at temperatures where *tost-1(xf196 ts)* alone supports normal development (at 15°C and 20°C) (Figure 3b). This finding demonstrates that TOST-1 and histone gene dosage interact genetically and is consistent with the idea that TOST-1 affects embryonic viability via maternal histone mRNA levels.

**Figure 3.**
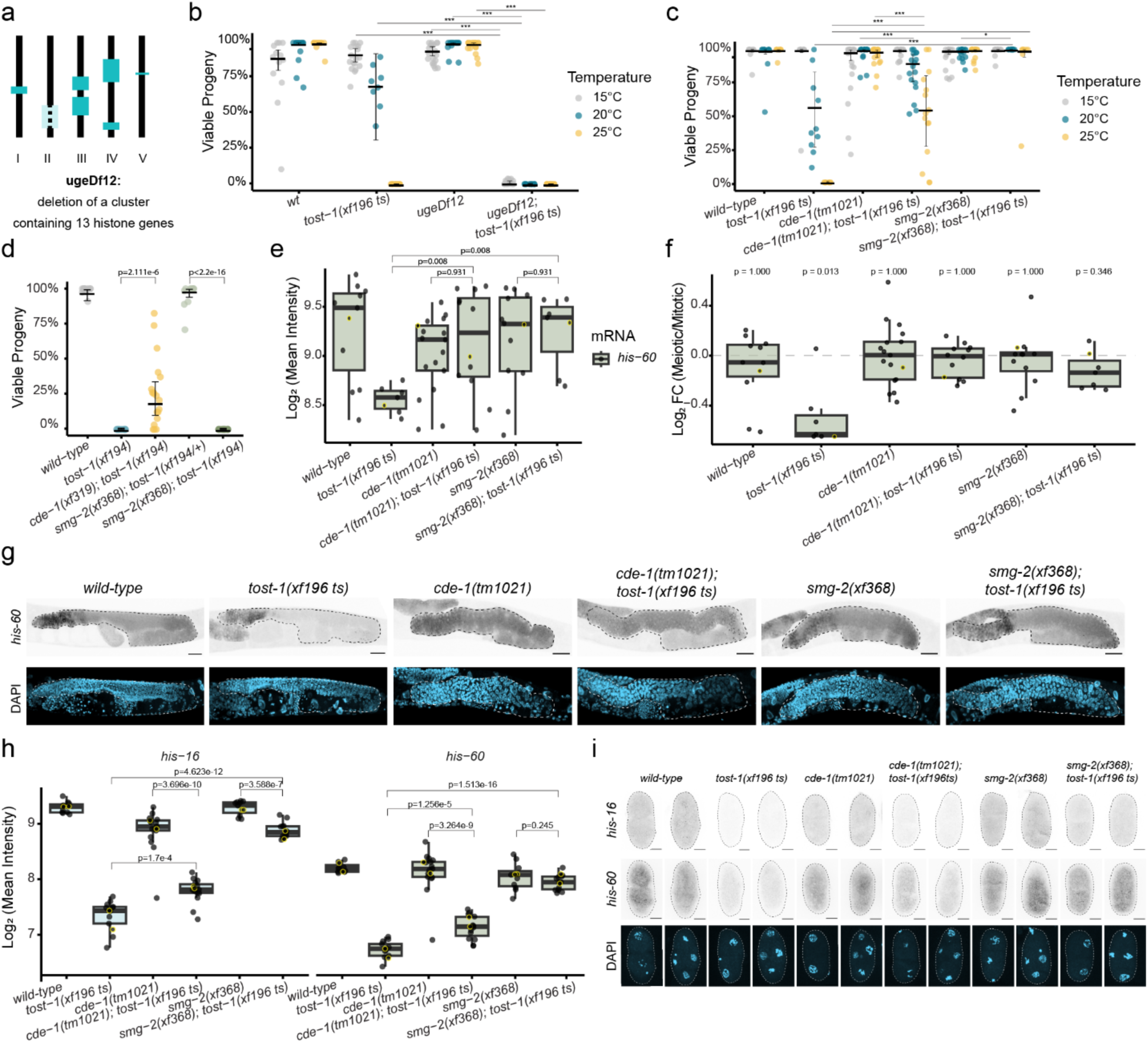
Genetic interactions between histone gene dosage and histone mRNA degradation pathways and *tost-1* mutants. **a,** Schematic representation of *C. elegans* chromosomes showing histone gene clusters and the region in chromosome II deleted in *ugeDf12*. **b,** Embryonic viability at 15°C, 20°C, and 25°C for *wild-type* (n=13, 17, 20 respectively), *tost-1(xf196 ts)* (n=18, 8, 16 respectively), *ugeDf12* (n=19, 17, 20 respectively), and *tost-1(xf196 ts); ugeDf12* double mutants (n=16, 11, 13 respectively). Statistical analysis using maximum a posteriori estimation within a Bayesian Generalised Linear Mixed-Effects Model with specific pairwise comparisons. Error bars represent 95% confidence intervals of the estimated viability. P values were corrected using the multivariate-t method within each temperature. Asterisks indicate significant differences: *P<0.05, **P<0.01, ***P<0.001. **c,** Embryonic viability at 15°C, 20°C, and 25°C of *wild-type* (n=9, 10, 9 respectively), *tost-1(xf196 ts)* (n=9, 10, 10 respectively), *cde-1(tm1021)* (n=20, 20, 18 respectively), *cde-1(tm1021); tost-1(xf196 ts)* (n=16, 17, 15 respectively)*, smg-2(xf368)* (n=19, 20, 19 respectively)*, and smg-2(xf368); tost-1(xf196 ts)* (n=16, 16, 8 respectively). Statistical analysis as in panel b. **b, c,** Exact P values are indicated in Extended Data Table 1. **d,** Embryonic viability at 20°C for *wild-type* (n=10), *tost-1(xf194)* (n=10), *cde-1(xf319); tost-1(xf194)* (n=19), *smg-2(xf368); tost-1(xf194/+)* (n=10), and *smg-2(xf368); tost-1(xf194)* (n=18). Statistical analysis as above. **e,** Quantification of *his-60* smFISH signal (log2 mean intensity) in adult gonads of *wild-type* (n=11), *tost-1(xf196 ts)* (n=7), *cde-1(tm1021)* (n=17), *cde-1(tm1021); tost-1(xf196 ts)* (n=12)*, smg-2(xf368)* (n=12)*, and smg-2(xf368); tost-1(xf196 ts)* (n=7). P values calculated using Welch’s t-test with Holm correction for multiple comparisons. **f,** Log2 fold change (FC) of *his-60* smFISH signal in meiotic versus mitotic regions from samples in panel e. P values calculated using paired t-test with Holm correction. **g,** Representative inverted fluorescence smFISH images corresponding to quantification in panel e. Scale bars, 20 μm. **h,** Quantification of *his-16* and *his-60* smFISH signal in embryos until 4-cell stage from *wild-type* (n=11), *tost-1(xf196 ts)* (n=13), *cde-1(tm1021)* (n=17), *cde-1(tm1021); tost-1(xf196 ts)* (n=15)*, smg-2(xf368)* (n=11)*, and smg-2(xf368); tost-1(xf196 ts)* (n=14). Statistical analysis as in panel e. **i,** Representative 2-cell and 4-cell stage embryo images corresponding to quantification in panel h. Scale bars, 10 μm. Yellow data points indicate samples corresponding to representative images shown.

### Histone mRNA decay pathway components suppress PETISCO phenotypes

To further test whether PETISCO functions through histone mRNA stabilisation, we investigated whether impairing histone mRNA degradation could rescue PETISCO mutant phenotypes. We examined mutations in *cde-1* (also known as *cid-1* and *pup-1*) and *smg-2*, the *C. elegans* homologs of TUT7 (a terminal uridylyl transferase) and UPF1 (an ATP-dependent RNA helicase), respectively, which participate in histone mRNA degradation in mammals^21,44^ but whose roles in nematode histone regulation are currently unknown.

Loss of CDE-1 rescued embryonic development in *tost-1* mutants: *cde-1* deletions restored viability in both the hypomorphic *tost-1(xf196 ts)* at restrictive temperature and remarkably, even in the null *tost-1(xf194)* allele (Figure 3c,d). Similarly, *smg-2* deletion completely rescued *tost-1(xf196 ts)* embryonic lethality, though it could not suppress the null allele phenotype (Figure 3c,d). This suggests that while *smg-2* loss can partially restore histone mRNA stability, this rescue is insufficient to compensate for the complete loss of TOST-1 function in the null mutant. This rescue extended across PETISCO components, with *cde-1* deletions restoring embryonic development in *pid-3, tofu-6* and *erh-2* mutants (Extended Data Fig 4). We note, however, that these rescue effects were not fully penetrant and that embryonic lethality tended to reappear in later generations, for reasons we do not yet understand.

SmFISH confirmed that both *cde-1* and *smg-2* mutations significantly rescued histone mRNA expression levels, restoring them back to the single mutant levels in adult germlines and reducing the characteristic difference of expression observed in meiotic regions of PETISCO mutants (Figure 3e-g). In embryos, histone expression levels were also rescued to different extents by loss of CDE-1 or SMG-2 (Figure 3h,i). We conclude that PETISCO and the histone degradation machinery operate in opposing functions to control histone mRNA homeostasis.

### A forward genetic screen identifies novel histone regulatory components

To discover additional factors in the PETISCO-histone mRNA pathway, we performed an unbiased EMS mutagenesis screen for suppressors of the *tost-1(xf196 ts)* Mel phenotype. L4 larvae of the mutant strain were mutagenized, and the F2 progeny were screened at the restrictive temperature (25°C) for viable individuals, indicative of suppression of the Mel phenotype. After backcrossing to the original mutant line, we recovered 21 suppressor lines that reproducibly rescued the Mel phenotype at 25°C (Extended Data Fig 5a). Whole-genome sequencing of these lines was performed using the original *tost-1(xf196 ts)* strain as reference for variant calling and linkage analysis (Extended Data Fig 5b).

This revealed mutations in three genes: *smg-1* (3 alleles), *smg-3* (1 allele), and *C14C10.5* (4 alleles) (Figures 4a-c and Extended Data Fig 5b). SMG-1 and SMG-3 encode components of the nonsense-mediated decay (NMD) pathway, functioning upstream of SMG-2/UPF1 in mRNA surveillance^45,46^. The *smg-1* mutations included a splice site disruption and lesions in the FAT and PI3K catalytic domains, all predicted to compromise kinase function (Figure 4a). The identified *smg-3* allele was predicted to truncate the protein immediately before a helix in the C-terminal region (Figure 4b). For both *smg-1* and *smg-3,* we generated deletion alleles using CRISPR-Cas9, which also rescued the *tost-1(xf196 ts)* Mel phenotype at 25°C (Figures 4d,e). This shows that besides SMG-2, also the NMD factors SMG-1 and SMG-3 have a role in histone mRNA homeostasis.

**Figure 4.**
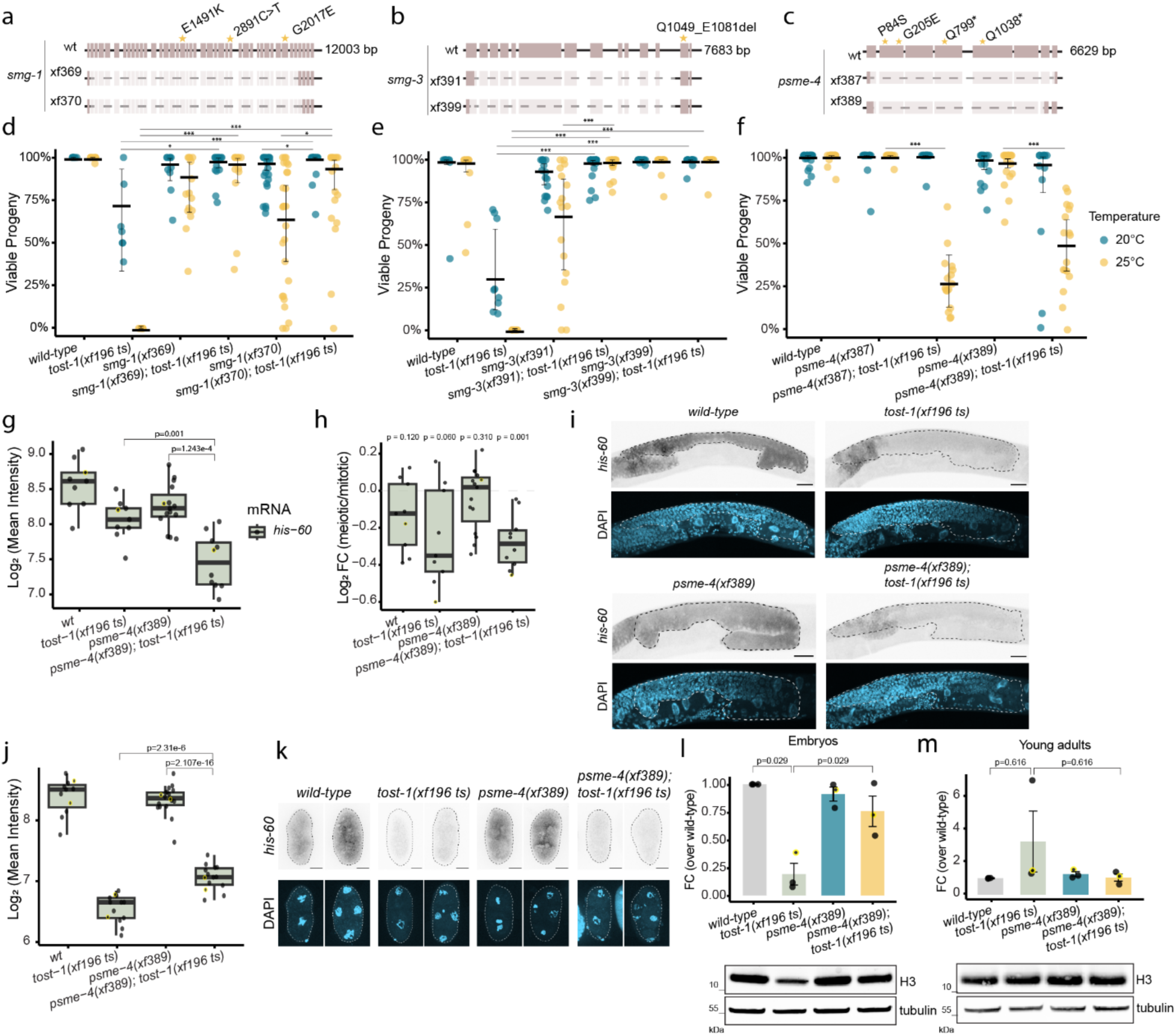
Forward genetic screen identifies histone mRNA and protein regulatory components. **a-c,** Schematic representation of *smg-1* (**a**), *smg-3* (**b**), and *psme-4 / C14C10.5* (**c**) genes showing EMS-induced mutations (indicated with a star) and CRISPR-Cas9 generated deletion alleles (light shaded regions). **d,** Embryonic viability at 20°C, and 25°C for *wild-type* (n=10, 10 respectively), *tost-1(xf196 ts)* (n=7, 7 respectively), *smg-1(xf361)* (n=10, 15 respectively), *smg-1(xf361); tost-1(xf196 ts)* (n=15, 12 respectively)*, smg-1(xf370)* (n=22, 26 respectively), and *smg-1(xf370); tost-1(xf196 ts)* (n=19, 18 respectively)*. **e,*** Embryonic viability at 20°C, and 25°C for *wild-type* (n=10, 10 respectively), *tost-1(xf196 ts)* (n=9, 8 respectively), *smg-3(xf391)* (n=16, 17 respectively), *smg-3(xf391); tost-1(xf196 ts)* (n=19, 16 respectively)*, smg-3(xf399)* (n=19, 18 respectively), and *smg-3(xf399); tost-1(xf196 ts)* (n=17, 16 respectively)*. **f,*** Embryonic viability at 20°C, and 25°C for *wild-type* (n=29, 29 respectively), *psme-4(xf387)* (n=19, 19 respectively), *psme-4(xf387); tost-1(xf196 ts)* (n=19, 16 respectively)*, psme-4(xf389)* (n=16, 20 respectively), and *psme-4(xf389); tost-1(xf196 ts)* (n=13, 17 respectively). **d-f,** Statistical analysis using maximum a posteriori estimation within a Bayesian Generalised Linear Mixed-Effects Model with multivariate-t multiple test correction within each temperature. Error bars represent 95% confidence intervals of the estimated viability. Exact P values are indicated in Extended Data Table 1. **g,** Quantification of *his-60* smFISH signal in adult gonads of *wild-type* (n=9), *tost-1(xf196 ts)* (n=9), *psme-4(xf389)* (n=15), and *psme-4(xf389); tost-1(xf196 ts)* (n=10) animals. P values calculated using Welch’s t-test with Holm correction. **h,** Log2 fold change (FC) of *his-60* smFISH signal in meiotic versus mitotic regions from samples in panel g. P values calculated using paired t-test with Holm correction. **i,** Representative inverted fluorescence smFISH images corresponding to quantification in panel g. Scale bars, 20 μm. **j,** Quantification of *his-60* signal in embryos until 4-cell stage from wild-type (n=11), *tost-1(xf196 ts)* (n=13), *psme-4(xf389)* (n=19), and *psme-4(xf389); tost-1(xf196 ts)* (n=14). Statistical analysis as in panel g. **k,** Representative embryo images corresponding to quantification in panel j. **l,** Western blot quantification of histone H3 protein levels in young adults (n=3 biological replicates per genotype). Expression normalised to tubulin and calculated as fold change (FC) relative to *wild-type*. P values calculated using one-sample t-test for *tost-1* mutant and paired t-test with Holm correction for comparison between *tost-1* mutant and *psme-4; tost-1* double mutant. Error bars represent SEM. Representative blot shown below. **m,** Western blot quantification of histone H3 protein levels in embryos (n=3 biological replicates per genotype). Analysis as in panel l. Yellow data points indicate samples corresponding to representative images shown.

The *C14C10.5* alleles included two premature stop codons and two missense mutations in exon 2, within an uncharacterised region of the protein (Figure 4c). Two deletion alleles of *C14C10.5,* made using CRISPR–Cas9, also restored viability to maternal *tost-1(xf196 ts)* embryos at 25°C, showing that *C14C10.5* is a *bona fide* suppressor of the *tost-1* Mel phenotype (Figure 4f). *C14C10.5* encodes the proteasome activator PSME-4 (human PSME4/PA200). In humans, this factor specifically targets acetylated histones for degradation during spermatogenesis and DNA repair^47^. Thus, we hypothesised that histone mRNA levels may not be restored in *psme-4; tost-1* double mutants, but that the rescue would act at the protein level. Consistent with this, histone mRNA levels remained reduced in *psme-4; tost-1* young adults and embryos. Additionally, smFISH analysis showed still a reduction in *his-60* signal in the meiotic versus mitotic regions (Figure 4g-k). Notably, *psme-4; tost-1* double mutant gonads show an even stronger reduction in *his-60* expression, possibly indicative of negative feedback mechanisms (Figure 4g). Yet, in *psme-4; tost-1* double mutant, histone H3 protein levels were higher compared with *tost-1(xf196 ts)* single mutant embryos (Figure 4l). In young adults, histone H3 protein levels did not significantly differ from controls, likely because histone H3 protein levels are not affected by TOST-1 in the adult in the first place (Figure 4m).

This demonstrates that the Mel phenotype resulting from PETISCO:TOST-1 dysfunction can be rescued by stabilising either histone mRNAs or proteins, highlighting the critical importance of maintaining adequate histone protein levels during early development.

### PETISCO directly binds replication-dependent histone mRNAs

To identify how PETISCO:TOST-1 affects histone mRNAs, we performed individual-nucleotide resolution UV crosslinking and immunoprecipitation (iCLIP) using an anti-TOFU-6 antibody in embryos (Extended Data Fig 6a,b). This analysis revealed highly specific enrichment of TOFU-6 binding in RD histone transcripts compared to controls, while most replication-independent (RI) histones showed no significant enrichment (Figure 5a). This specificity correlates with the presence of a conserved stem-loop structure that distinguishes RD from RI histone mRNAs. Beyond the annotated RD histones, TOFU-6 also bound *his-39*, a histone H2B variant, and *his-69*, a histone H3.3, although both retain the stem-loop structure despite being classified as RI. A metagene analysis of crosslinked sites, as defined by the position upstream of the 5’ ends of the mapped reads, across all RD histone transcripts revealed that TOFU-6 binds preferentially in the 3’ regions (Figure 5b), with a strong binding peak occurring precisely 12 nucleotides upstream of the stem-loop structure (Figures 5b,c and Extended Data Fig 6c-e). This position corresponds to the start of a conserved sequence motif present in all RD histone genes in *C. elegans*, but not, for instance, in human (Figure 5e), suggesting that PETISCO recognises a specific cis-regulatory element. In addition, analysis of the 3’ ends of the bound transcripts indicated that PETISCO bound predominantly to mature, processed histone mRNAs rather than to longer precursors (Figure 5d). We conclude that PETISCO:TOST-1 mainly binds to mature RD histone mRNAs and that it contacts a conserved sequence element just upstream of the 3’ end stem-loop.

**Figure 5.**
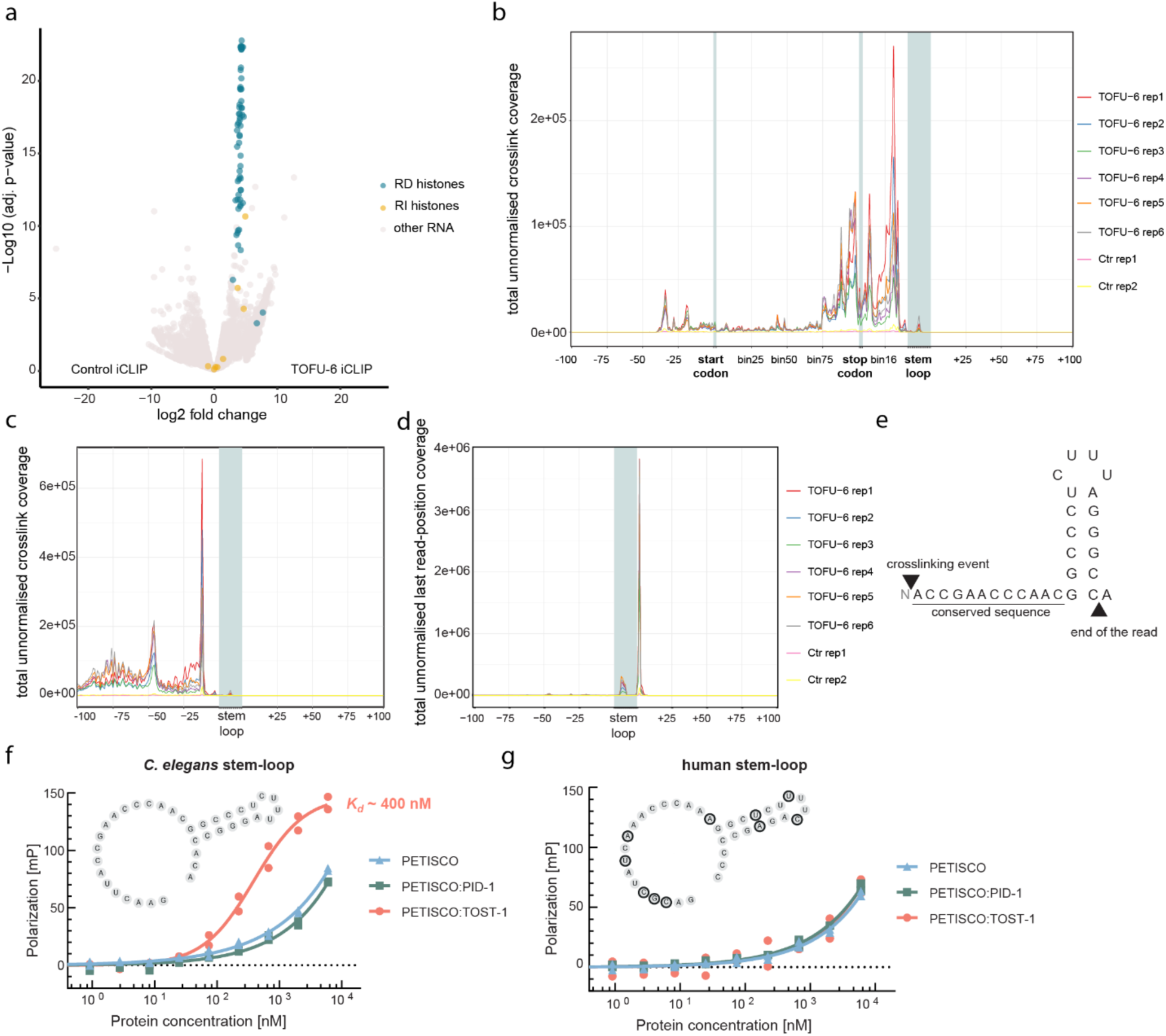
PETISCO binds RD histone mRNAs via the stem-loop in a TOST-1-dependent manner. **a,** Volcano plot showing the results of a comparison of TOFU-6 iCLIP (n=6) and control iCLIP (n=2) to identify significant TOFU-6 binding. Genes are plotted by binding fold change (log2) versus statistical significance (-log10(adjusted P value)). Histone genes are labelled as replication-dependent (RD) or replication-independent (RI). **b,** Meta-coverage plot showing the total coverage of crosslink sites (first position upstream of each read) across all RD histone transcripts. The coding sequence (CDS) between start and stop codons was divided into 100 bins adjusting for gene length differences between genes; similarly the region between stop codon and stem-loop was divided into 31 bins. **c,** Meta coverage plot showing the total coverage of crosslink sites relative to the stem-loop region across all RD histone genes per TOFU-6 iCLIP replicate. **d,** Meta-coverage plot showing total coverage of read termination sites of mapped trimmed reads (last position of reads) relative to the stem-loop region of RD histone genes. **b-d**, Data are shown unnormalised because the control libraries contained very low amounts of material, leading to low read numbers. Normalisation would artificially inflate control profiles to inappropriate levels. **e,** Schematic of canonical RD histone stem-loop structure showing upstream conserved sequence motif across *C. elegans* RD histone genes. Arrowheads indicate predominant read start and end positions relative to the stem-loop. **f,** Fluorescence polarisation assays measuring binding of PETISCO alone, PETISCO:PID-1, and PETISCO:TOST-1 to fluorescently labelled *C. elegans* histone stem-loop RNA (n=2 replicates each). Stem-loop structure of RNA substrate shown. **g,** Fluorescence polarisation assays as in panel f using labelled human histone stem-loop RNA (n=2 replicates each). Stem-loop structure shown with nucleotides differing from *C. elegans* sequence highlighted in blue.

To probe the interaction of the histone stem-loop region with recombinant proteins, we purified PETISCO in three different forms: bound to TOST-1, bound to PID-1 and not bound to either of these two proteins (Extended Data Fig 6f). Then, we used a fluorescence polarisation assay to probe the binding specificities of these complexes: PETISCO:TOST-1 bound efficiently (Kd≈400nM) to *C. elegans* histone sequences containing the upstream motif and stem-loop from *his-8*, but showed much weaker binding to the equivalent human histone sequences or control RNAs (Figure 5f,g and Extended Data Fig 6g,h). Efficient binding was strictly dependent on TOST-1 presence, establishing that the PETISCO:TOST-1 complex, not PETISCO alone nor PETISCO:PID-1, can mediate histone mRNA recognition. Notably, RD histone mRNAs in *C. elegans* lack the tri-methyl-G-capped trans-spliced leader sequences found on most nematode transcripts (Extended Data Fig 6i); as a result they retain a 5’ m^7^G cap structure. This unique feature may contribute to their recognition by the 5’ m^7^G cap-binding protein IFE-3 within PETISCO, providing an additional layer of specificity for histone mRNA binding. These findings establish that PETISCO:TOST-1 functions as a sequence-specific histone mRNA-binding complex, directly recognising and stabilising the maternal histone transcripts essential for early embryonic development.

## Discussion

Precise regulation of histone abundance is fundamental to genome stability and developmental timing across eukaryotes^13,14,48,49^. Perturbations in maternal histone supply have severe consequences during embryogenesis, as observed in flies, fish and frogs. When histone levels are excessive, transcription is delayed and nuclear divisions increase, conversely, histone depletion accelerates ZGA onset, lengthens cell cycles, and triggers checkpoint responses^5–7^. In addition, defective chromosome condensation, resulting in developmental arrest, has been observed in histone-depleted cells^34,35,50–52^. All these processes rely on the protein SLBP.

RD histone transcription is typically restricted to S phase to ensure stoichiometric histone supply during DNA replication^14,53,54^. However, exceptions exist during oogenesis when large maternal stores of both histone mRNAs and proteins accumulate and are stabilised for use during embryogenesis^17,19,26^. In frogs, histone mRNAs made during oogenesis are stored in association with an oocyte-specific SLBP, which represses translation^17,28^. Similarly, zebrafish employ an oocyte-specific SLBP that is also important for regulating histone mRNA and protein storage during oogenesis^30^. In *Drosophila*, histone genes are transcribed in stage 10B nurse cells independent of DNA replication, with mRNAs deposited into oocytes where both histone mRNA and protein are stored^24,26^. Recent work found that a pool of maternal histone mRNAs is polyadenylated with truncated stem-loops, revealing an SLBP-dependent process that likely enhances transcript stability during storage^55^. This polyadenylation of maternal RD histone mRNAs is conserved in *Xenopus*^28,56^, and polyadenylated histone mRNAs have also been reported in *C. elegans*^57^. Our findings do not exclude the possibility that a similar polyadenylation mechanism is relevant to maternal histone mRNAs in nematodes. Interestingly, mouse oocytes maintain histone mRNA stability for some histone genes even when SLBP fails to accumulate in mature oocytes^19,52^, suggesting that SLBP-independent mechanisms may exist also in mammals.

PETISCO likely stabilises histone mRNAs while maintaining translational repression, a function consistent with the role of its IFE-3 subunit, which is an eIF4E homolog^58,59^. Consistent with our data, IFE-3 RIP experiments detected a clear enrichment of RD histone mRNAs^60^. Curiously, a zebrafish eIF4E homolog, eIF4E1b binds to maternal short polyadenylated transcripts, including non-polyadenylated histone mRNAs, to prevent premature translation^31^. By sequestering histone transcripts in the gonad, PETISCO could simultaneously protect them from degradation and block untimely translation, functionally replacing the poly(A)-tail-mediated repression that is commonly used for other maternal mRNAs^61,62^. Mechanistically, PETISCO’s described associations with Y-box proteins (CEY proteins in *C. elegans*) and the DEAD-box helicase CGH-1^37,38,63^, both involved in maternal mRNA dormancy^64–66^, suggest that PETISCO integrates into previously described RNA regulatory networks to ensure maternal histone mRNAs remain available for use during embryogenesis.

A question that awaits further experimentation is how PETISCO-bound RD histone mRNAs can be used in the early embryos. Possibly, the mere availability of CDL-1/SLBP could prime the release of the mRNA from PETISCO, based on differences in affinity: SLBP has been reported to have a Kd of roughly 1nM towards the histone stem-loop^67^ whereas PETISCO:TOST-1 binds more than two orders of magnitude weaker to this region of the histone mRNA. We note, however, that the additional presence of the m^7^Cap and binding to IFE-3 would most likely increase the affinity of PETISCO. In this light, the spatial separation of CDL-1 in oocyte nuclei in *C. elegan*s^68^, away from the cytoplasmic TOST-1^38^, may be a prerequisite for the system to function. The regulation of CDL-1 localization or activity in oocytes and early embryos may depend on post-translational modifications. For instance, it was demonstrated previously that SLBP phosphorylation affects its stability and stem-loop binding affinity^69,70^.

In certain PETISCO mutants, especially the *tost-1* hypomorphic animals, we observed RD histone mRNAs accumulating in foci in and/or around the nuclei. Notably, this was most evident in the region of the gonad, roughly the diplotene region, where wild-type animals displayed an increase in histone mRNA levels. These foci may relate to their transcription, their decay, to storage or any combination of these options. A role in transcription should involve foci within the nuclei. Unfortunately, however the resolution of the fixed smFISH samples prevents us from making clear statements on whether we see intranuclear foci or not. Roles in either decay or storage could involve P bodies. Indeed, PETISCO components have been detected in P-bodies and other germ granule types^60,71^, and a single allele of *patr-1*, the *C. elegans* homolog of the human P-body component PATL1/2, was identified in our genetic screen, but not yet validated. These findings suggest that at least part of the identified foci may be P-bodies, related to the decay of histone mRNAs.

Another striking insight is the evolutionary connection between histone mRNA regulation and piRNA biogenesis. PETISCO’s dual role, stabilising histone transcripts when bound to TOST-1 and piRNA precursors when bound to PID-1^36,37^, illustrates how molecular platforms can be repurposed across pathways. Both histone mRNAs and piRNAs undergo specialised 3′-end processing distinct from canonical mRNAs, and unlike these canonical mRNAs, they are not spliced. Moreover, it has been suggested that both share common transcription termination factors, such as the integrator complex^72,73^. The recent discovery of a common AT-hook transcription factor regulating histone mRNAs, snRNAs and SL RNAs in *C. elegans* further supports this convergence^74^. TOST-1’s deeper conservation compared to PID-1^37^, implies that histone mRNA regulation represents PETISCO’s ancestral function, with piRNA biogenesis layered onto this machinery later during nematode evolution.

Taken together, we have identified a unique SLBP-independent mechanism for regulating RD histone mRNA, ensuring stability and translational repression outside S phase. PETISCO’s modularity, achieved by a single alternative effector protein, illustrates an elegant evolutionary principle: complex regulatory capacity emerges not through *de novo* invention, but through the tweaking and repurposing of existing modules.

## Materials and Methods

### 1. Worm culture

*C. elegans* strains were maintained at 20°C on NGM plates seeded with *E. coli* OP50 according to standard laboratory conditions^75^. MosSCI strains were cultured on plates seeded with *Comamonas* sp. (DA1877). For iCLIP, animals were grown on high-density OP50 egg plates for one generation^76^, synchronised by hypochlorite treatment, and embryos harvested in the next generation. For IP-MS, animals were grown on 90 mm NGM plates for one generation, synchronised, then transferred to 150 mm plates for one additional generation before harvest. Bristol N2 served as wild-type control. A list of the strains used is provided in Supplementary Table 1. Many aspects of this work made use of WormBase^77^.

### 2. MosSCI transgenesis

The MosSCI system was used to introduce transgenes into the *ttTi5605* locus on chromosome II^78,79^. For *xfSi254* transgene, the *gfp* coding sequence with three introns was amplified from pDD282 (Addgene #66823). For *xfSi268* transgene, the GFP coding sequence (without introns) was synthesised as a gBlock (IDT). Plasmids were purified using NucleoSpin® Plasmid kits (Macherey-Nagel®) and verified by PCR and sequencing. DNA mixes containing pCFJ601 (50 ng/μl), pMA122 (10 ng/μl), pGH8 (10 ng/μl), pCFJ104 (5 ng/μl), pCFJ90 (2.5 ng/μl), and either pRK3105 (*xfSi254*) or pRK3118 (*xfSi268*) (50 ng/μl each) were injected into both gonads of 50 young adult EG6699 worms. Progeny were screened for transgene integration as described^78,79^. Insertion events were confirmed by sequencing, and strains were outcrossed twice before analysis. A list of plasmids used is provided in Supplementary Table 5.

### 3. CRISPR/Cas9 genome editing

Protospacer sequences were selected using CRISPOR^80^ and cloned into pRK2411 (derived from pDD162, Addgene #47549) via site-directed, ligase-independent mutagenesis (SLIM)^81^. SLIM reactions were transformed into DH5α competent cells (Invitrogen™) and plated on LB agar with 100 μg/ml ampicillin. Genome editing used *dpy-10(cn64)* or *unc-58(e665)* co-conversion strategies^82^. Protospacer sequences and repair templates are listed in Supplementary Table 3 and 4, respectively. DNA mixes were injected into both gonad arms of 10-20 young adult N2 hermaphrodites at 20°C. F1 progeny were screened by PCR and confirmed by Sanger sequencing. Generated mutant strains were outcrossed at least twice before analysis. A list of the genotyping primers is provided in Supplementary Table 6.

### 4. Embryonic viability assay

Mutant strains were confirmed and outcrossed twice before experiments. Worms were synchronised by hypochlorite treatment and hatched in M9 buffer for 16 h. Synchronised L1 larvae were grown to L4 stage, then individual worms were transferred to 60 mm NGM plates at appropriate temperatures (15°C, 20°C, or 25°C). After 24-48h reproduction, adult progenitors were removed and embryos counted. Plates were incubated for an additional 48-72h before recording hatched larvae. Embryonic viability was calculated as the proportion of hatched embryo to total embryos. A filter of minimum of 10 embryos was used in these experiments, with the exception in the Extended Data Fig 1b, were this filter was eliminated due to the low broad size of the animals.

### 5. RNAi treatment

RNAi experiments were conducted as described^83,84^. For *C. briggsae*, RNAi-competent strain JU1018 was used. A 100-200 bp exon sequence from each target gene was PCR-amplified using primers with T7 promoter sequences. PCR products were cloned into L4440 plasmid by Gibson assembly^85^ and transformed into HT115 *E. coli*. Exon sequences are listed in Supplementary Table 8. For *C. elegans,* the *tost-1, tofu-6* and *his-65* targeting RNAi constructs were obtained from the Ahringer RNAi library^86,87^. NGM plates supplemented with 1mM IPTG were seeded with transformed bacteria to induce dsRNA synthesis.

### 6. Widefield microscopy

Adult hermaphrodites were washed and dissected in M9 buffer before mounting on glass slides with 2% agarose pads. Transmitted and fluorescence imaging of embryos was performed using Leica DM6000B or DMi8 widefield (Thunder, Leica Microsystems) with LED excitation, respective filter set for GFP, a 63×/1.43 oil immersion objective and sCMOS camera (Flash V4.2, Hamamatsu or K7, Leica Microsystems) detection, controlled by LASX (v3.7) software. Images were processed and analysed using Fiji^88^ and Omero software^89,90^.

### 7. Live imaging

#### a. Sample preparation and imaging

Adult hermaphrodites were dissected in M9 buffer to isolate embryos and mounted as described. Time-lapse imaging was performed over 66 minutes with 3-minute intervals at 20°C. Images were acquired with a spinning disk confocal microscope (VisiSope 5Elements) based on a Nikon Ti-2E stand, Yokogawa CSU-W spinning disk (50 μm pinhole), and controlled by VisiView® software. The system was equipped with a 60x water immersion objective (CFI Plan Apo VC 60XWI, 1.2 NA), 2x magnification lens, a Prime BSI sCMOS camera (2048 x 2048 pixels, 6.5 µm pixel size, Photometrics), and with an incubator (Bold Line universal stage top incubator, okolab). Embryos were excited at 488 nm (argon laser, 20% power) with emission detected at 500-550 nm. Z-stacks of whole samples were acquired with 0.5 μm step size. Raw images (.nd files) stored in Omero software^89,90^.

#### b. Image processing and quantification

For time-lapse experiments, z-stacks were combined using sum of slices Z-projection in Fiji^88^. GFP mean intensity and standard deviation were calculated across entire embryos after background subtraction, with regions of interest (ROIs) manually defined using the brightfield channel. For nuclear analysis, average GFP mean intensity was calculated from three in-focus nuclei per embryo. When nuclear GFP expression was absent, ROIs were defined using brightfield.

### 8. RNA extraction and RNA sequencing

Synchronised gravid adults were washed with M9 buffer and subjected to hypochlorite treatment to isolate embryos. Embryos were washed four times with cold M9 buffer and frozen as “worm balls” in liquid nitrogen. Frozen samples were ground with mortar and pestle, then mixed with five volumes of TRIzol™ LS Reagent (Invitrogen™). RNA was extracted using Direct-zol RNA Microprep kit (Zymo Research™) with additional TURBO DNase treatment (Invitrogen™) to remove genomic DNA. RNA was resuspended in nuclease-free water.

#### a. Library preparation and sequencing

NGS libraries were prepared using Illumina’s Stranded mRNA Prep Ligation Kit with 1000 ng starting material and 10 amplification cycles. ERCC spike-ins (2 μl of 1:100 dilution, Ambion) were added to assess technical variability and optionally assist with library size normalization. Libraries were profiled using DNA 1000 chip on 2100 Bioanalyzer (Agilent) and quantified with Qubit dsDNA HS Assay Kit. All samples were pooled equimolarly and sequenced on NextSeq 500 (1×80 cycles plus 10 index cycles).

#### b. Read processing and differential expression analysis

Sample demultiplexing used bcl2fastq (v2.19.1.403). Raw reads were quality-assessed with FastQC (v0.11.8) and aligned to the *C. elegans* genome (WBcel235/ce11 assembly) with gene annotation from Ensembl release 104 using STAR (v2.7) with parameter “--outFilterMismatchNoverLmax 0.04”. The genome FASTA file and the annotation GTF file were supplemented with data of the added ERCC spike-ins. Secondary alignments were removed with Samtools (v1.9). Data quality was assessed using QualiMap (v2.2.1) and dupRadar (v1.37.0). Read counts were summarised on the gene level with Subread featureCounts (v1.6) using stranded parameter “-s 2” and the above mentioned GTF annotation file. Differential expression analysis was performed using DESeq2 (v1.26.0) following the standard data processing workflow with the default median-ratio normalisation, dispersion estimation, negative binomial GLM fitting and Wald statistics for pairwise comparisons with independent gene filtering and 1% FDR threshold.

#### c. Gene expression fraction during embryogenesis

Identifiers of genes with significant changes to mRNAs were extracted from R along with the top 25% genes with highest baseMean in the DESeq analysis and uploaded to GExplore^40,91^, from where their life stage specific expression profiles were downloaded. All genes were normalised to their own expression to generate expression fractions relative to developmental progression and embryo time point expression fractions were plotted with ggplot2 (v3.5.1) in R. The shaded area is the confidence interval of the geom_smooth function.

#### d. Gene Ontology enrichment analysis

GO enrichment analysis was performed on differentially expressed genes using the clusterProfiler (v4.6.2) package in R with the *C. elegans* annotation database (org.Ce.eg.db). Upregulated and downregulated genes were analysed separately against DESeq2 tested genes as background. Enrichment was tested across all GO categories with p-value cutoff of 0.05 and adjusted using the Benjamini-Hochberg method.

### 9. smFISH

#### a. Sample preparation and imaging

smFISH probes targeting *his-16* and *his-60,* labelled with Quasar570 and Quasar670 dyes respectively were designed using Stellaris Probe Designer (Supplementary Table 3). For larvae, 60 animals were washed in M9 buffer and fixed in 4% paraformaldehyde in 1× PBS for 1 h at room temperature. After centrifugation and PBS wash, worms were permeabilised overnight in 70% ethanol at 4°C. For embryos, samples were harvested three days after hypochlorite treatment for synchronization and fixed in 4% PFA in 1× PBS for 15 min at room temperature. Samples were vortexed, freeze-cracked in liquid nitrogen for 1 min, thawed at room temperature, vortexed again, incubated on ice for 20 min, and washed twice with PBS. Embryos were permeabilised overnight in 70% ethanol at 4°C. Hybridization was performed with probes at 125 nM in hybridization buffer (100 mg/ml dextran sulfate, 1 mg/ml *E. coli* tRNA, 2 mM vanadyl ribonucleoside complex, 0.2 mg/ml RNase-free BSA, 10% formamide) overnight at 30°C. Samples were washed twice for 30 min at 37°C with wash buffer (10% formamide, 1× SSC), with DAPI (5 ng/ml) added to the final wash. Embryos received a final 5-min wash with 2× SSC at room temperature, while larvae were washed with 2× SSC, PBS + 1% Tween, and PBS. Samples were mounted in glycerol based mounting medium (Ibidi) and imaged using a Spinning Disc Confocal Microscope BC43 (Oxford Instruments, Andor), controlled by Fusion software with a 40×/0.75 air objective or 100×/1.45 oil objective. Z-stacks covering the full sample were acquired with 0.4 μm steps size. Raw images (.ims files) stored in Omero software^89,90^.

#### b. Image processing and analysis

Fiji was used for smFISH image analysis based on z-projection intensity (sum of slices)^88^. For larval samples, the gonadal region of interest (ROI) was segmented using the *his-60* smFISH (Quasar670) or GFP::WAGO-3 signals, while the entire animal was segmented using the DAPI channel. The background area was defined as the region outside the whole-animal ROI. Within each gonad, mitotic and meiotic regions were manually defined based on DAPI morphology: the mitotic ROI extended from the distal tip to the transition zone (identified by the first crescent-shaped nuclei), and the meiotic ROI extended from the transition zone to the last oocyte. For embryo samples, the *his-60* smFISH signal was used to define embryo ROIs, with the inverse regions used as background. All ROIs were applied to the blue (DAPI), red (*his-16* smFISH), and far-red (*his-60* smFISH) channels to extract area, mean, standard deviation, modal, minimum, maximum, and integrated density values. Signal intensities were background-corrected by subtracting the mean background value from each corresponding ROI measurement.

### 10. Quantitative RT-qPCR

RNA samples were prepared as described above. Reverse transcription was performed using normalised total RNA with ProtoScript First Strand cDNA Synthesis Kit (NEB) and qPCR Random Primer Mix. qPCR reactions (10 μl) contained PowerUp™ SYBR™ Green Master Mix (Applied Biosystems), 500 nM primers, and cDNA diluted to ∼1 ng/μl. Amplifications were performed on Applied Biosystems ViiA7 Real Time PCR System using standard cycling conditions: 95°C for 10 min, followed by 40 cycles of 95°C for 15 s and 60°C for 1 min, with melt curve analysis. Technical triplicates and biological duplicates or triplicates were used as indicated. Data were analysed using the ΔΔCT method^92^ with *pmp-3* as reference gene^93^. Error bars represent 95% confidence intervals of log2 fold changes. Primers are listed in Supplementary Table 7.

### 11. Ems screening

#### a. EMS mutagenesis

Mixed-stage plates of RFK912 strain containing early L4 larvae were washed with M9 buffer and collected in 15 ml tubes. Worm pellets were resuspended in 3 ml M9 buffer and transferred to tubes containing EMS (Sigma) diluted to 47 mM final concentration. Worms were incubated at 20°C for 4 h on a spinning wheel, then washed twice with M9 buffer and plated on NGM plates at 15°C. After seven days, 1100 F1 offspring were singled (5 per 60 mm plate) at 15°C. When F1 reached L4 stage, plates were transferred to 25°C. After five days, plates were screened for F3 suppressors, which were singled and maintained at 25°C for one generation to confirm homozygosity. F4 animals with viable progeny were backcrossed twice to RFK912 strain and expanded for genomic DNA extraction.

#### b. Library preparation

Genomic DNA was extracted using Gentra Puregene Tissue Kit (Qiagen). DNA (1.5 μg) was diluted to 55 μl in TE buffer and fragmented using Covaris S2 sonicator (Intensity: 5; Duty Cycle: 10%; Cycles/burst: 200; Time: 120 s; 2 cycles). Fragmented DNA was analysed on TapeStation with High Sensitivity D1000 ScreenTape. Libraries were prepared using NEBNext Ultra II DNA Library Prep Kit starting with 10 ng fragmented DNA and 9 PCR cycles. Libraries were profiled on 2100 Bioanalyzer (Agilent) and quantified using Qubit 1× dsDNA HS Assay Kit. All 22 samples were pooled equimolarly.

#### c. Next-generation sequencing data analysis

Libraries were sequenced on NextSeq 2000 (100 bp paired-end). Reads were adapter-trimmed using Cutadapt (v4.4)^94^ and mapped to *C. elegans* genome (WBcel235) using BWA-MEM2 (v2.2.1)^95^. Duplicates were removed with Picard (v3.0.0)^96^ and tracks generated using bamCoverage (v3.5.1)^97^. Variants were called using GATK HaplotypeCaller (v4.4.0.0)^98^ and filtered for SNPs, indels, and mixed variants. Additional filtering included variants present in RFK912 (*tost-1(xf196 ts))* sample (background control), standard quality filters, and minimum depth of 6 using vcftools (v0.1.16)^99^. Variants were classified as homozygous/heterozygous using GATK tools, converted to MAF format, and annotated using VEP (v110.1). Expected G→A and C→T transitions were selected using bcftools (v1.17), and filtered VCF files were converted to XLSX format using R/Bioconductor packages.

### 12. Western blot

For adult samples, 100 non-gravid hermaphrodites were hand-picked into 1× NuPAGE™ LDS sample buffer (Invitrogen™) supplemented with 100 mM DTT and incubated at 95°C for 30 min. For embryo samples, 200 gravid adults were hand-picked in M9 buffer, subjected to hypochlorite treatment to dissolve adults while preserving embryos, washed three times with M9 buffer, then processed with LDS sample buffer as above. Following mixing and centrifugation, supernatants were transferred to fresh tubes and stored at −20°C. Samples were separated on NuPAGE™ 10% Bis-Tris gels in MES SDS running buffer at 120 V alongside Color Prestained Protein Standard (10–250 kDa, NEB). Proteins were transferred to Protan BA85 nitrocellulose membrane (Amersham) at 120 V for 1 h using NuPAGE™ Transfer Buffer with 10% methanol. Membranes were blocked in PBS containing 5% skim milk and 0.1% Tween-20 for 1 h, then sectioned by molecular weight. Sections were incubated overnight at 4°C with primary antibodies: anti-TOFU-6 (in-house), anti-α-tubulin (mouse, 1:1000, Abcam ab7291), or anti-H3 (rabbit, 1:5000, Sigma-Aldrich H0164) in blocking buffer. After three 10-min washes with PBS + 0.1% Tween, membranes were incubated 1 h with LI-COR secondary antibody IRDye 800CW (goat, 1:10000, LI-COR Biosciences 926-32211) and washed three times. Protein detection was performed using LI-COR Odyssey M Western blot imager.

### 13. Anti-PID-3 and anti-TOFU-6 antibody production

Plasmid pET28a carrying 6×His-TOFU-6 or PID-3 were transformed into BL21(DE3) cells (Thermo Fisher) and grown overnight in LB with kanamycin at 37°C. Culture was diluted 1:100 in 2 L total volume, grown to OD₆₀₀ = 0.8, cooled on ice for 15 min, and induced with 1 mM IPTG. Cells were grown at 16°C, harvested by centrifugation (4000×g, 20 min), and resuspended in PBS with benzonase (1:5000). Cells were disrupted using Cell Disruptor TS2 at 4°C and centrifuged (19,500×g, 30 min). The pellet was washed in cold IMAC A buffer (PBS, 8 M urea, 20 mM imidazole), resuspended in 20 ml IMAC A, and incubated 1 h on ice with vortexing. After sonication and centrifugation (4000×g, 30 min), supernatant was loaded onto 5 ml HisTrap column (Cytiva) using ÄKTA Prime at 1 ml/min. The column was washed with 50 ml IMAC A and eluted with gradient IMAC B (50 mM Tris-HCl pH 8.0, 300 mM NaCl, 10% glycerol, 500 mM imidazole, 1 mM DTT). Fractions were analysed by SDS-PAGE. Urea concentration was reduced to 4 M during purification, and purified proteins were used for antibody production. Antibodies were raised in rabbit (Eurogentec).

#### a. Antibody affinity purification

Recombinant PID-3 and TOFU-6 were dialysed into coupling buffer (50 mM Tris pH 8.5, 150 mM NaCl, 5 mM EDTA, 6 M Urea, 1 mM TCEP) for 2h at room temperature. 2 mg of each protein was covalently coupled to equilibrated SulfoLink resin (Thermo Fisher Scientific) at 1 mg/ml for 2 h at room temperature in Poly-Prep columns (Bio-Rad) while rotating. After washing with coupling buffer, the remaining iodoacetyl groups were blocked with quenching buffer (50 mM L-cysteine in coupling buffer) for 30 min. The resin was subsequently washed with coupling buffer containing 1 M NaCl, followed by PBS. For antibody purification, rabbit serum (2-10 ml) was filtered through 0.45 μm filters and applied to antigen-coupled SulfoLink resin in Poly-Prep columns overnight at 4°C. Resin was washed sequentially with PBS, PBS containing 0.1% Triton X-100, and PBS. Antibodies were eluted with glycine buffer (200 mM glycine-HCl, 150 mM NaCl, pH 2.3) and immediately neutralised with 1 M Tris pH 8.5. Eluted fractions were analysed by SDS-PAGE, rebuffered into storage buffer (PBS, 10% glycerol, 0.05% NaN3) using PD-10 columns (Cytiva). Antibodies were concentrated to 1 µg/µl using Amicon Ultra-4 spin-concentrators with 10 kDa cut-off (Merck) and aliquots were flash-frozen and stored at -80°C.

### 14. iCLIP

#### a. Samples and library preparation and sequencing

iCLIP in *C. elegans* embryos was performed as described^100^. Wild-type worms were grown on egg plates and embryos harvested by hypochlorite treatment. Embryos were UV cross-linked four times at 100 mJ/cm² (254 nm) in Worm Lysis Buffer (25 mM Tris-HCl pH 7.5, 150 mM NaCl, 1.5 mM MgCl₂, 1 mM DTT, 0.1% Triton X-100) and snap-frozen as “worm balls.” DYNAL™ Dynabeads™ Protein G (100 μl) were conjugated with 2 μg TOFU-6 antibody for 1 h at room temperature and washed twice. Embryo extracts (1 mg total protein per replicate) were treated with RNase I and TURBO DNase at 37°C for 3 min, clarified by centrifugation, and filtered. Lysates were incubated with antibody-conjugated beads for 2 h at 4°C, washed, and treated with T4 PNK for RNA 3’-end dephosphorylation (20 min, 37°C). L3-App adapter ligation was performed overnight at 16°C, followed by RNA 5’-end labelling with ³²P-γ-ATP (37°C, 5 min). Samples were run on 4-12% NuPAGE Bis-Tris gels, transferred to nitrocellulose membrane, and visualised by phosphoimaging. RNA-protein complexes were isolated, treated with Proteinase K, and RNA extracted using phenol/chloroform. After precipitation, reverse transcription was performed using SuperScript III, followed by second adapter ligation and cDNA amplification (18-24 cycles). Libraries were assessed using TapeStation and quantified by Qubit. Six replicates and two negative controls (without antibody) were pooled equimolarly and sequenced on an Illumina NextSeq 500 sequencing machine as 150 nt single-end reads including a 6 nt sample barcode as well as 5+4 nt unique molecular identifiers (UMIs).

#### b. Sequencing and read processing and analysis

Basic quality controls were done with FastQC (v0.11.9) (https://www.bioinformatics.babraham.ac.uk/projects/fastqc/) and reads were filtered based on sequencing qualities (Phred score) in the barcode and UMI regions using the FASTX-Toolkit (v0.0.14) (http://hannonlab.cshl.edu/fastx_toolkit/) and seqtk (v1.3) (https://github.com/lh3/seqtk/). Reads with a Phred score below 10 in the considered regions were removed from further analysis. The remaining reads were de-multiplexed based on the sample barcode on positions 6 to 11 of the reads using Flexbar (v3.5.0)^101^. Barcode and UMI regions as well as adapter sequences were trimmed from read ends using Flexbar requiring a minimal overlap of 1 nt of read and adapter. UMIs were added to the read names and reads shorter than 15 nt were removed from further analysis.

Duplicated reads were defined as identical reads including identical UMIs. They were removed from the de-multiplexed and trimmed reads of each sample using basic Bash commands. De-duplicated reads were mapped using STAR (v2.7.3a)^102^ and genome assembly WBcel235 and annotation of Ensembl release 108^103^. During mapping, up to 4% of the bases were allowed to be mismatched (--outFilterMismatchNoverReadLmax 0.04 --outFilterMismatchNmax 999), a splice junction overhang of 134 nt (--sjdbOverhang 134) was used and soft-clipping at the 5’ end (--alignEndsType Extend5pOfRead1) was prohibited. Since many reads were expected to map to multiple locations, the maximally accepted number of locations was set to 999 (--outFilterMultimapNmax 999). Multiple alignments of reads were output in random order and the choice of the primary alignment from the highest scoring alignments was made at random (--outMultimapperOrder Random). Secondary hits were removed using Samtools (v1.10)^104^ (samtools view -F 256) keeping only one hit per multi-mapping read. The resulting BAM files were sorted using Samtools (v1.10). Reads directly mapped to the chromosome ends were removed since they do not have an upstream position and, thus, no crosslink position can be extracted.

The position upstream of mapped reads was extracted using BEDTools (v2.29.2)^105^ bamtobed, shift (with parameters -m 1 -p -1) and genomecov (with parameters -bg -strand + -5 for reads on the forward strand and with parameters -bg -strand - -5 for reads on the reverse strand). The resulting bedGraph files were converted to bigWig files using bedGraphToBigWig of the UCSC tool suite (v385)^106^. In addition to the position upstream of mapped reads, the last position of mapped reads, which were trimmed at the 3’ end by at least 6 nt, was extracted as well using BEDTools (v2.29.2) bamtobed and genomecov (with parameters -bg -strand + -3 for reads on the forward strand and with parameters -bg -strand - -3 for reads on the reverse strand). Requiring at least 6 nt to be trimmed ensures that the last position of the reads is indeed the end of the RNA insert. The resulting bedGraph files were converted to bigWig files using bedGraphToBigWig of the UCSC tool suite (v385).

Exonic reads per gene were counted using featureCounts from the Subread tool suite (v2.0.0)^107^ with non-default parameters --donotsort -M -s1. Genes significantly bound by TOFU-6 were identified by comparing TOFU-6 iCLIP and control iCLIP samples using functions of the R package DESeq2 (v1.38.1)^108^ with a false discovery rate (FDR) threshold of 1% in an R/Bioconductor environment (v3.16/v4.2.2)^109,110^. As control iCLIP samples only differ from TOFU-6 iCLIP samples by not using the TOFU-6 antibody, we are confident that the differences we see are due to different binding behaviour and not due to different expression. However, to be sure, this would require expression data in addition to the iCLIP data to be confirmed. Results from DESeq2 were visualised as a volcano plot showing the log2 fold change of normalised counts of TOFU-6 iCLIP and control iCLIP conditions versus the negative log10 of adjusted P values (corrected for multiple testing using the method of Benjamini and Hochberg). RD histone genes are highlighted in blue, while RI histone genes are shown in yellow. Histone genes were divided between RD and RI based on the presence of the conserved stem-loop, except for *his-69* and *his-39* that although are also considered RI, have a stem-loop^35^.

Using bigWig coverage tracks of either positions upstream of mapped reads or last positions of mapped trimmed reads, we extracted the coverage +/-100 nt around the start and stop codons as well as around the stem-loop regions of RD histone genes using R packages rtracklayer (v1.58.0)^111^, GenomicRanges (v1.50.1)^112^, stringr (v1.5.0)^113^, Biostrings (v2.66.0)^114^, and BSgenome (v1.66.1)^115^ in an R/Bioconductor environment (v4.2.2/v3.16)^109,110^. Start and stop codon locations were taken from WBcel235 annotation of Ensembl release 108^103^. In addition to extracting coverages around start and stop codons as well as stem-loop regions separately, we extracted the coverage of the complete gene starting 100 nt upstream of the start codon and ending 100 nt downstream of the stem-loop region using bigWig coverage tracks of positions upstream of mapped reads and R packages rtracklayer (v1.58.0), GenomicRanges (v1.50.1), stringr (v1.5.0), Biostrings (v2.66.0), and BSgenome (v1.66.1) in an R/Bioconductor environment (v4.2.2/v3.16). Start and stop-codons as well as stem-loop regions were kept as anchor points when comparing different genes. Since genes have different lengths (CDS region) and also different distances between the stop codon and the stem-loop sequence, these regions have to be made equally long in all genes when they are summarised and shown in the same plot. This was done by equal binning in these regions, i.e. the same number of bins was used for every gene. Since the region between start and stop-codon is typically several 100 nt long, this region was split into 100 equal sized bins and the average position-wise coverage per bin was calculated. The region between the stop codon and the stem-loop sequence varies between 31 and 57 nt. It was split into 31 bins for all considered genes and the average position-wise coverage per bin was calculated. The regions upstream of the start codon as well as downstream of the stem-loop sequence were not binned, but kept as 100 individual positions. The total coverage of all RD histone genes was calculated per sample.

### 15. Recombinant protein production

PETISCO, PETISCO:PID-1, and PETISCO:TOST-1 complexes were assembled from N-terminally His-tagged fusion proteins expressed in *E. coli* BL21(DE3) derivatives grown in terrific broth medium. Cultures were grown at 37°C to OD₆₀₀ = 2, cooled to 18°C for 2 h, induced with 0.2 mM IPTG, and incubated overnight (12-16 h) at 18°C. Cell pellets were mixed in specific ratios: TOFU-6:IFE-3 heterodimer with PID-3:ERH-2 tetramer (2:3 wet weight) for PETISCO, or with PID-3:ERH-2:PID-1 or PID-3:ERH-2:TOST-1 hexamers (1:2 ratio). Pellets were resuspended in lysis buffer (25 mM Tris-HCl pH 8.0, 50 mM sodium phosphate, 500 mM NaCl, 20 mM imidazole, 10% glycerol, 5 mM 2-mercaptoethanol, 1 mM PMSF, 14 nM benzonase) and lysed by sonication. EDTA-free cOmplete protease inhibitor (Roche) was added for PETISCO:PID-1. Complexes were purified by IMAC using 5 ml Ni²⁺-chelating HisTrap FF columns (Cytiva) and eluted with 25 mM Tris-HCl pH 8.0, 50 mM sodium phosphate, 500 mM NaCl, 500 mM imidazole, 10% glycerol, 5 mM 2-mercaptoethanol. His-tags were removed by 3C protease during overnight dialysis against 25 mM Tris-HCl pH 8.0, 150 mM NaCl, 5% glycerol, 5 mM 2-mercaptoethanol. Further purification used 5 ml HiTrap Heparin HP columns followed by size-exclusion chromatography on HiLoad 16/600 Superdex 200 column in 20 mM Tris-HCl pH 7.5, 150 mM NaCl, 5% glycerol, 2 mM DTT. All steps were performed at 4°C.

### 16. RNA production

RNAs were produced by *in vitro* transcription using homemade T7 polymerase and partially complementary ssDNA templates (Supplementary Table 9). Transcription reactions contained 40 mM Tris-HCl pH 8.0, 10 mM each rNTP, 50 mM MgCl₂, 1 mM spermidine, 5 mM DTT, 0.005% IGEPAL CA-630, 0.1 U/μl RNasin (Promega), 0.1 U/μl pyrophosphatase (Thermo), 2.5 μM pre-mixed primers, and 20 μg/ml T7 polymerase. Reactions were incubated for 6 h at 37°C with shaking. DNA templates were digested with 0.005 U/μl DNase I (Thermo) plus 0.5 mM CaCl₂ for 1 h at 37°C. Reactions were stopped with 100 mM EDTA and incubated at 75°C for 15 min. RNAs were precipitated overnight at -20°C using 300 mM sodium acetate and 0.7 volumes of isopropanol. RNAs were purified by anion exchange chromatography using HiTrap Capto DEAE columns (Cytiva) and treated with homemade RppH to convert the 5’-triphosphate to monophosphate. For 3’-end fluorescent labelling, 200 μg RNA was oxidized with 100 mM sodium meta periodate in 100 mM sodium acetate (4 h, 4°C). The remaining oxidation reagent was precipitated with 180 mM KCl. The RNA was labelled with 6 mM fluorescein-5-thiosemicarbazide (FTSC, Sigma Aldrich) overnight at 4°C. Excess dye was removed by phenol/chloroform/isoamyl alcohol extraction, and RNAs were precipitated and desalted using NAP-5 columns (Cytiva).

### 17. Fluorescence polarization assays

A constant concentration of RNA (10 nM) was titrated with increasing concentrations of protein complexes in 180 μl total volume. Buffer contained 20 mM Tris-HCl pH 7.5, 150 mM NaCl, 1 mM EDTA, 2 mM DTT, and 0.005% IGEPAL CA-630. After 10 min incubation at room temperature, fluorescence polarization was measured on a Tecan Spark plate reader (excitation 485 nm ± 20 nm, emission 510 nm ± 10 nm). Polarization values (mP) were normalised by subtracting background from wells containing only labelled RNA. Data were analysed using nonlinear regression with the Hill equation in GraphPad Prism (v10.2.3). Experiments were performed in duplicate with individual replicates plotted.

### 18. Immunoprecipitation

Synchronised *C. briggsae* were cultivated at 20°C, harvested at gravid-adult stage, and frozen in four 200 μl aliquots in sterile water on dry ice. Thawed samples were combined with equal volume of 2× lysis buffer (50 mM Tris-HCl pH 7.5, 300 mM NaCl, 3 mM MgCl₂, 2 mM DTT, 0.2% Triton X-100, cOmplete Mini EDTA-free protease inhibitors). Samples were sonicated using Bioruptor Plus (4°C, 10 cycles, 30 s ON/30 s OFF), centrifuged (21,000×g, 10 min, 4°C), and protein concentrations quantified by BCA assay. Samples were split and diluted to 1.5 mg in 400 μl lysis buffer for PID-3 IP and IgG control. For each IP, 30 μl DYNAL Dynabeads Protein G were conjugated with 2 μg PID-3 antibody or anti-IgG control antibody (CST). Beads were washed three times with wash buffer (25 mM Tris-HCl pH 7.5, 150 mM NaCl, 1.5 mM MgCl₂, 1 mM DTT, protease inhibitors) and pre-incubated with worm extract (1-2 h, 4°C). Pre-cleared extracts were combined with antibody-conjugated beads and rotated for 2 h at 4°C. Beads were washed three times and resuspended in 25 μl 1.2× NuPAGE LDS sample buffer with 120 mM DTT.

### 19. Mass Spectrometry

#### a. Sample preparation and analysis

Immunoprecipitation experiments were performed in quadruplicate as described above. Eluted proteins were processed using SP3 approach^116^, digested with trypsin overnight at 37°C, and purified using C18 StageTips.

#### b. Liquid chromatography tandem mass spectrometry

Peptides were separated on an in-house packed 30-cm analytical column (75 μm inner diameter, ReproSil-Pur 120 C18-AQ 1.9-μm beads, heated to 50°C) using 105-min non-linear gradient of 1.6-32% acetonitrile with 0.1% formic acid at 225 nl/min. Eluted peptides were analysed by Q Exactive Plus Orbitrap mass spectrometer (Thermo Scientific) in data-dependent acquisition mode using top10 method: one full scan (300-1,650 m/z, resolution 70,000, target 3×10⁶, 20 ms injection time) followed by 10 HCD fragmentation scans (25% normalised collision energy, resolution 17,500, target 1×10⁵, 120 ms injection time, 1.8 m/z isolation window). Precursor ions of unassigned or +1 charge were rejected; selected ions were excluded for 20 s.

#### c. Data processing and statistical analysis

Raw data were processed using MaxQuant (v2.1.3.0) with Andromeda search engine against target-decoy database containing UniProt *C. briggsae* (release 2023_02, 21,756 entries), *E. coli* (release 2023_01, 5,064 entries), and common contaminants. Parameters: trypsin/P specificity, carbamidomethylation of cysteine (fixed), methionine oxidation and N-terminal acetylation (variable), maximum 2 missed cleavages, 1% FDR at peptide and protein levels. MaxLFQ algorithm performed label-free quantification without default normalization, minimum ratio count of 1. *E. coli* proteins, reverse hits, contaminants, and “only identified by site” groups were filtered out. Data were log-transformed and median-centrered. Proteins detected in ≥2/4 replicates per group were retained. Statistical significance assessed using modified t-statistic (SAM) with thresholds t₀ = 1.4 and s₀ = 1.5.

### 20. Yeast Two-hybrid

Genes of interest were amplified from cDNA and cloned into pGAD and pGBD plasmids^117^. List of plasmids present in Supplementary Table 5. All plasmids were transformed into haploid *Saccharomyces cerevisiae* strain AH109 using high-efficiency LiAC method^118^ with denatured salmon sperm DNA (95°C, 5 min). Transformants were cultured on agar plates supplemented with adenine (ADE) and histidine (HIS). For interaction selection, transformants were plated at 5×10⁶ and 5×10⁵ cells/ml on plates lacking HIS. Plates contained Yeast Synthetic Drop-out Medium Supplements without histidine, leucine, tryptophan, and adenine (1.399 g/L) and Yeast Nitrogen Base Without Amino Acids (6.7 g/L). Selection plates were supplemented with adenine (21 mg/L) and L-histidine monohydrochloride (85.6 mg/L), plus 20 g/L agar and 2% glucose.

### 21. 5’ Rapid Amplification of cDNA Ends (RACE) PCR

RNA was purified from mixed-stage wild-type worms and cDNA synthesised using ProtoScript First Strand cDNA Synthesis Kit (NEB). 5’ RACE was performed using FirstChoice® RLM-RACE Kit (Invitrogen™) following manufacturer’s instructions. Specific oligonucleotides targeting *his-67*, *his-46*, and *tost-1* 5’ UTRs were used to assess trans-splicing events (Supplementary Table 10). 5’ RACE products were cloned using TOPO™ TA Cloning™ kit and transformed into One Shot™ TOP10 competent cells (Invitrogen). Clones were confirmed by Sanger sequencing.

### 22. Statistical analysis and reproducibility

Statistical analyses were performed using R-based packages (v4.2.2). Embryonic viability data were analysed using Bayesian mixed-effects logistic models. These models provide more robust handling of issues arising from quasi-separated data compared to standard generalised mixed-effects models. Replicates were modelled as random effects. Replicates with <10 embryos were excluded, except otherwise stated. The model used binomial family with logit link, fitting genotype as fixed effect and replicate as random effect. Estimated marginal means and 95% confidence intervals were calculated using *emmeans*^119^ (v1.10.5) and transformed to probability scale. Pairwise comparisons used contrast analysis with Tukey or multivariate-t correction for multiple testing. Analysis performed in R using *blme*^120^ (v1.0-6) and *emmeans* packages. Exact P values are shown in Extended Data Table 1. All experiments were repeated at least once.

In smFISH experiments, log-transformed background-corrected intensities were used for statistical analysis to normalise distributions. For the whole gonad and embryo analysis, comparisons between genotypes used Welch’s t-test on log-transformed intensities. P values were adjusted using Holm correction for multiple comparisons across targets (*his-16* and *his-60*). For mitotic vs. meiotic region analysis, fold changes between meiotic and mitotic regions were calculated as the difference in log-transformed intensities (log[meiotic] - log[mitotic]) for each sample. Paired t-tests compared mitotic and meiotic intensities within each genotype. P values were adjusted using Holm correction for multiple comparisons.

In quantitative RT-qPCR experiments, statistical comparisons between genotypes used Welch’s t-test on ΔCt values at each temperature. Log2 fold changes were calculated as the difference between group means. P values were adjusted using Bonferroni correction within each temperature group. Confidence intervals were calculated from t-test results and used as error bars.

In the western-blots, histone H3 protein levels were normalised to tubulin and expressed as fold change relative to *wild-type*. Statistical comparisons used: (1) paired t-test comparing *tost-1* versus *tost-1; psme-4* double mutant, (2) one-sample t-test comparing *tost-1* fold change against theoretical mean of 1.0 (*wild-type* level). P values were adjusted using Holm correction for multiple comparisons. Error bars represent standard error of the mean.

In the assay to analyse the premature transgene expression, nuclear GFP intensity was background-corrected and log-transformed for analysis. Mean intensity was calculated from three nuclei per embryo at time points 4, 10, 16, and 22. Linear regression analysis was performed on log-transformed intensity as a function of time and RNAi treatment interaction for time points >4: lm(log_intensity ∼ time * RNAi_treatment). The model tested whether GFP expression onset differed between RNAi conditions by comparing regression slopes (rate of intensity increase over time) and intercepts (initial expression levels on t=0) across treatments. P-values from the regression coefficients assessed: (1) whether each RNAi treatment significantly altered the slope compared to control, indicating changes in transcriptional activation kinetics, and (2) whether intercepts differed between RNAi treatments, indicating shifts in basal expression timing. Detection limit was established as the mean log intensity of all samples at time point 4.

## Data Availability

All data supporting the findings of this study are provided in the manuscript and its supplementary information. RNA-seq datasets generated in this study have been deposited in ArrayExpress under accession code E-MTAB-15730. iCLIP datasets have been deposited in ArrayExpress under accession code E-MTAB-15713. Whole-genome sequencing data from EMS suppressor lines are available in ArrayExpress under accession code E-MTAB-15751. Proteomics data from the IP–MS experiments have been deposited in UCSD MassIVE under accession code MSV000099274. All microscopy images are available through a publicly accessible OMERO database at https://omero.imb.uni-mainz.de/pub/pereirinha2025. Additional data are available from the corresponding author upon reasonable request.

## Acknowledgements

We thank all members of the Ketting laboratory for all the discussions and critical reading of the manuscript. We thank Svenja Hellman for technical support, and Ricardo Cordeiro Rodrigues for his contribution to the early stages of this project. Carolina Ruivinho and Vindhya Jaya are acknowledged for their contributions during the mutant screen. We thank Fridolin Kielisch from the IMB Bioinformatics Core Facility for consulting on statistical methods. We thank Emil Karaulanov and Nastasja Kreim from the IMB Bioinformatics Core Facility for RNA-sequencing and whole-genome analysis. We acknowledge support from members of the IMB Genomics Core Facility and use of the NextSeq 500 system (funded by the DFG, INST #329045328). We thank Martin Möckel from the IMB Protein Production Core Facility for antibody purification and other consumables. We thank the IMB Proteomics Core Facility for sample preparation, measurement and data analysis; measurements were performed on the Q Exactive Plus (#240874965). We thank Sandra Ritz, Marton Gelleri and Petri Turunen from the IMB Microscopy Core Facility for support and assistance. Live cell imaging was performed on a spinning disk microscope (VisiScope, Visitron) funded by DFG INST #402386039) We thank the IMB Media Laboratory for buffers and worm plates. Some strains were provided by the *Caenorhabditis* Genetics Center (CGC), funded by NIH Office of Research Infrastructure Programs (P40 OD010440) and by the National BioResource Project (NBRP), funded by the Japanese government. We thank the Kemphues Lab for providing additional *C. elegans* strain. This project was funded by the Deutsche Forschungsgemeinschaft (DFG, German Research Foundation) – GRK2526 – Project no. 407023052, 439669440-TRR319, 252386272 and 504320275 (to R.F.K.) and the Austrian Science Fund (FWF) I6110-B (to S.F.).

## Author contributions

J.P. and R.F.K. conceived the study and designed experiments. J.P. executed experiments and performed data analysis. M.B. performed PETISCO protein purification and fluorescence polarization assays. S.G. performed the experiments with *C. briggsae*. A.B. performed iCLIP analysis. N.P. purified proteins for antibody production and validated PID-3 and TOFU-6 antibodies. A.S.S. performed computational analysis of RNA-seq data. K.D. and F.A.S. provided unpublished strains. J.K. assisted with iCLIP experiments. S.F. assisted with experimental design and supervised M.B. R.F.K. supervised the project. J.P. and R.F.K. wrote the manuscript with input from all authors.

---

**Extended Data Figure 1.**
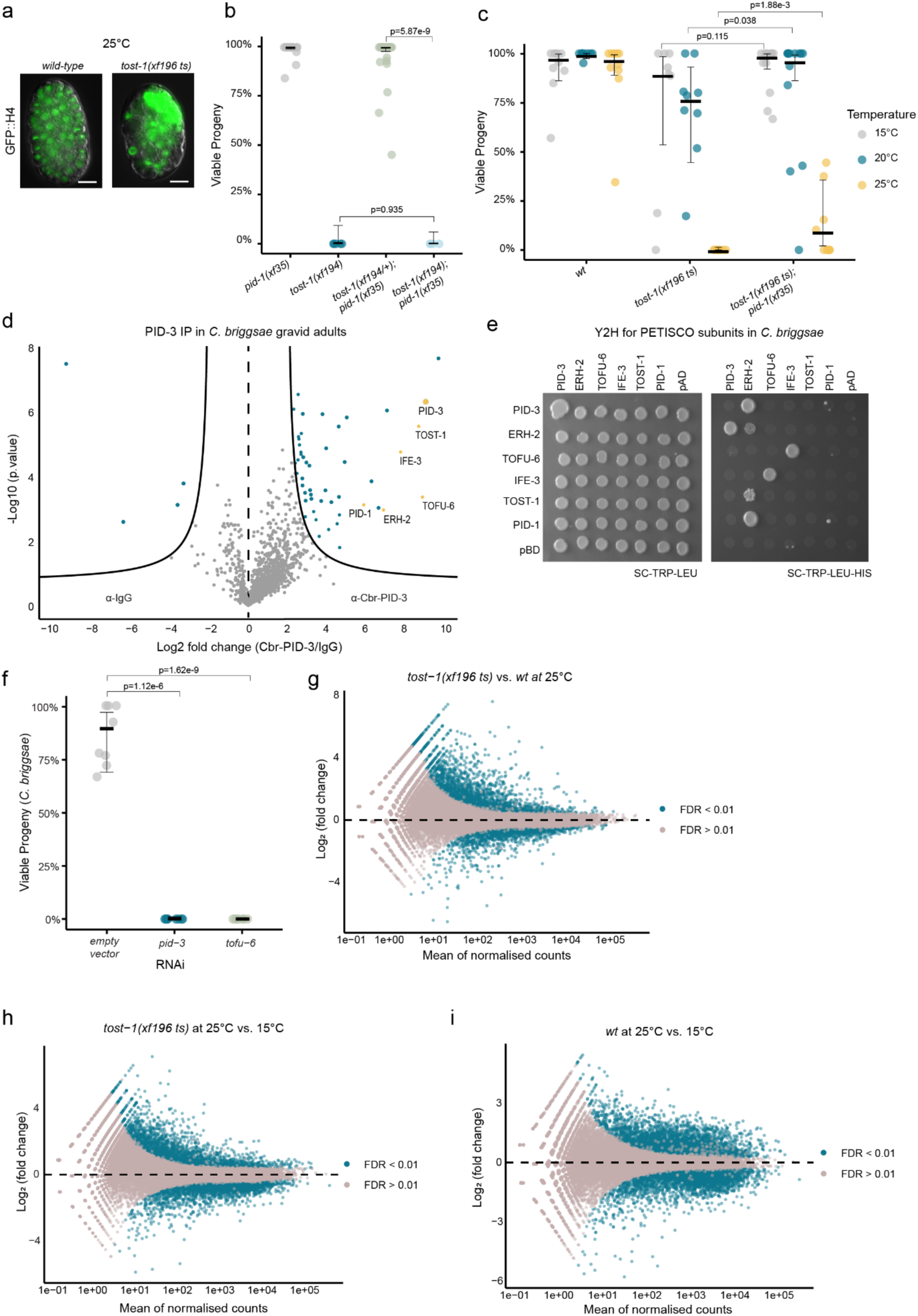
TOST-1 and PETISCO functions are evolutionarily conserved. **a,** Representative fluorescence images showing mitotic defects in *tost-1(xf196 ts)* mutants. Transgene *xfSi254*[GFP::H4] in *wild-type* (left) or *tost-1(xf196 ts)* (right) background grown at 25°C. Scale bar, 20 μm. **b,** Embryonic viability of *tost-1(xf194)* (n=5), *pid-1(xf35)* (n=18), *tost-1(xf194/+); pid-1(xf35)* (n=26), and *tost-1(xf194); pid-1(xf35)* (n=4) embryos. Statistical analysis using maximum a posteriori estimation within a Bayesian Generalised Linear Mixed-Effects Model with specific pairwise comparisons and multivariate-t method for multiple comparisons. Error bars represent 95% confidence intervals of the estimated viability. **c,** Embryonic viability at 15°C, 20°C, and 25°C of *wild-type* (n=10 in each temperature), *tost-1(xf196 ts)* (n=8, 9 and 9 respectively), and *tost-1(xf196 ts); pid-1(xf35)* (n=13, 13 and 9 respectively) mutants. Statistical analysis as in panel b. **d,** Immunoprecipitation-mass spectrometry in *C. briggsae* gravid adults using PID-3 antibody versus anti-IgG control. Lines show thresholds at P = 0.05 and twofold enrichment. Coloured data points represent proteins that meet both the threshold for statistical significance (t.sam = 1.4) and twofold enrichment. PETISCO components indicated in yellow. **e,** Yeast two-hybrid analysis of PETISCO subunit interactions. Left: control growth medium (SC without tryptophan and leucine); right: selective medium (SC without tryptophan, leucine, and histidine). **f,** Embryonic viability assay using RNAi with empty vector control (n=8), *pid-3* RNAi (n=8), or *tofu-6* RNAi (n=7). Statistical analysis as in previous viability assays. **g-i,** Differential expression MA plot of RNA-seq analysis of **(g)** *tost-1(xf196 ts)* vs. *wild-type* embryos at 25°C, **(h)** *tost-1(xf196 ts)* at 25°C vs. 15°C, **(i)** *wild-type* at 25°C vs. 15°C. Blue dots indicate significantly upregulated and downregulated genes (FDR <0.01). Adjusted P values calculated using Benjamini-Hochberg method. n = 3 biological replicates.

**Extended Data Figure 2.**
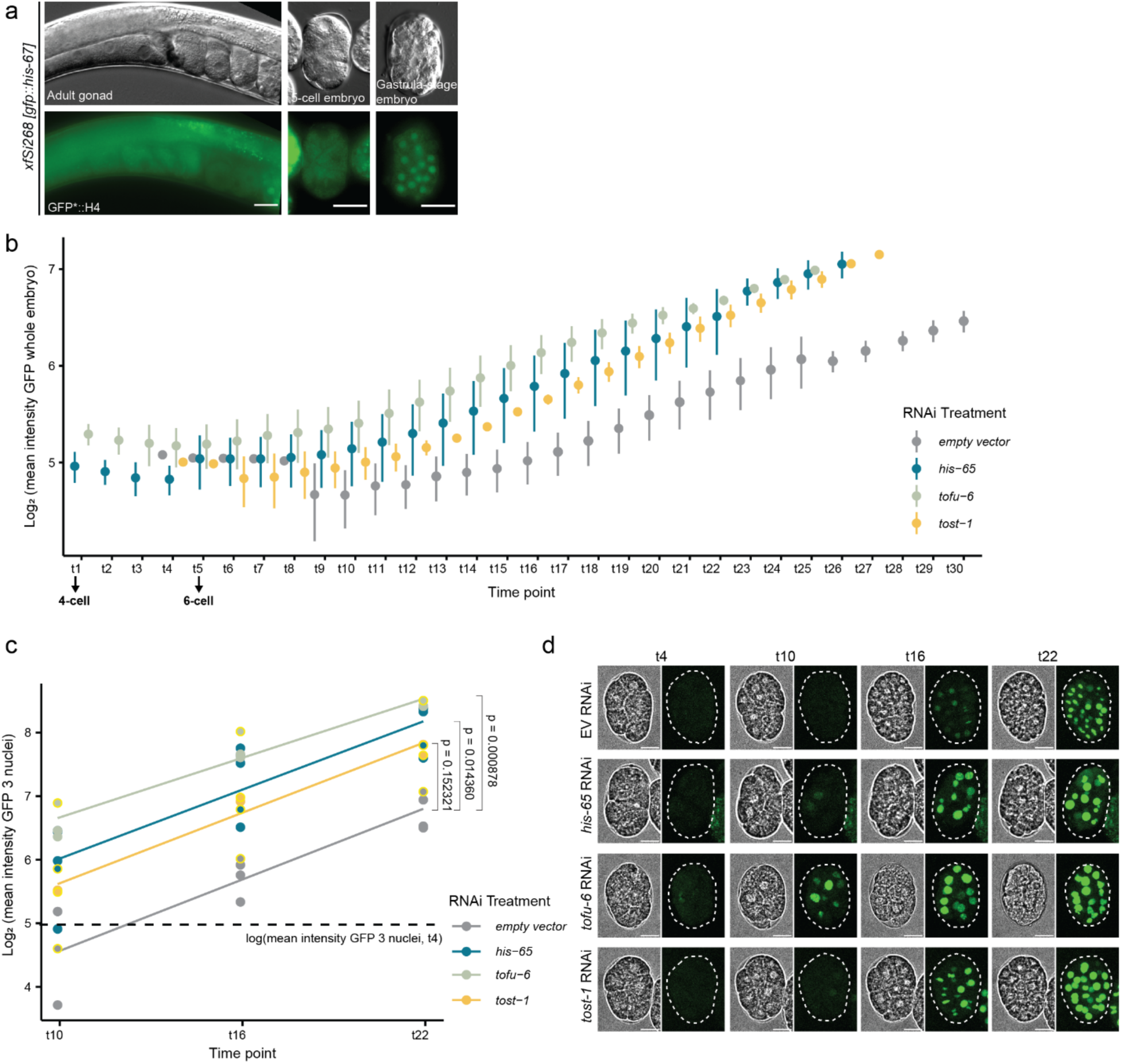
TOST-1 and PETISCO depletion advances zygotic transcription timing. **a,** Wide-field micrographs showing single optical sections of adult gonads and dissected embryos containing *xfSi268* transgene expressing GFP::H4 (top: DIC contrast, below: fluorescence contrast). Scale bars = 20 μm. **b,** Average fluorescence mean intensity of GFP::H4 expression during time-lapse spinning disk confocal imaging with 3-minute intervals. Embryos dissected from adults fed RNAi for empty vector (n=4), *his-65* (n=5), *tofu-6* (n=3), *tost-1* (n=2). GFP mean intensity measured in whole embryos, normalised by cell number. Time points adjusted with time point 1 as 4-cell embryo and time point 5 as 6-cell embryo. Error bars represent standard deviation across biological replicates. **c,** Linear regression analysis of nuclear GFP intensity over time. Mean intensity measured in 3 nuclei per embryo at time points 10, 16, and 22 for animals treated with empty vector RNAi, *his-65*, *tofu-6*, and *tost-1*. Scatter plot shows log intensity values from individual samples across time points. Lines represent model predictions from linear regression of log intensity as a function of time with interaction by RNAi treatment. Dashed horizontal line indicates mean log intensity at time point 4, used as detection limit. P values shown indicate significance of intercept differences (earlier expression onset) compared to control; slope differences were not significant (exact P values in Extended Data Table 1). **d,** Representative brightfield and z-projected fluorescence images of samples highlighted in yellow in panel c. Scale bars, 10 μm.

**Extended Data Figure 3.**
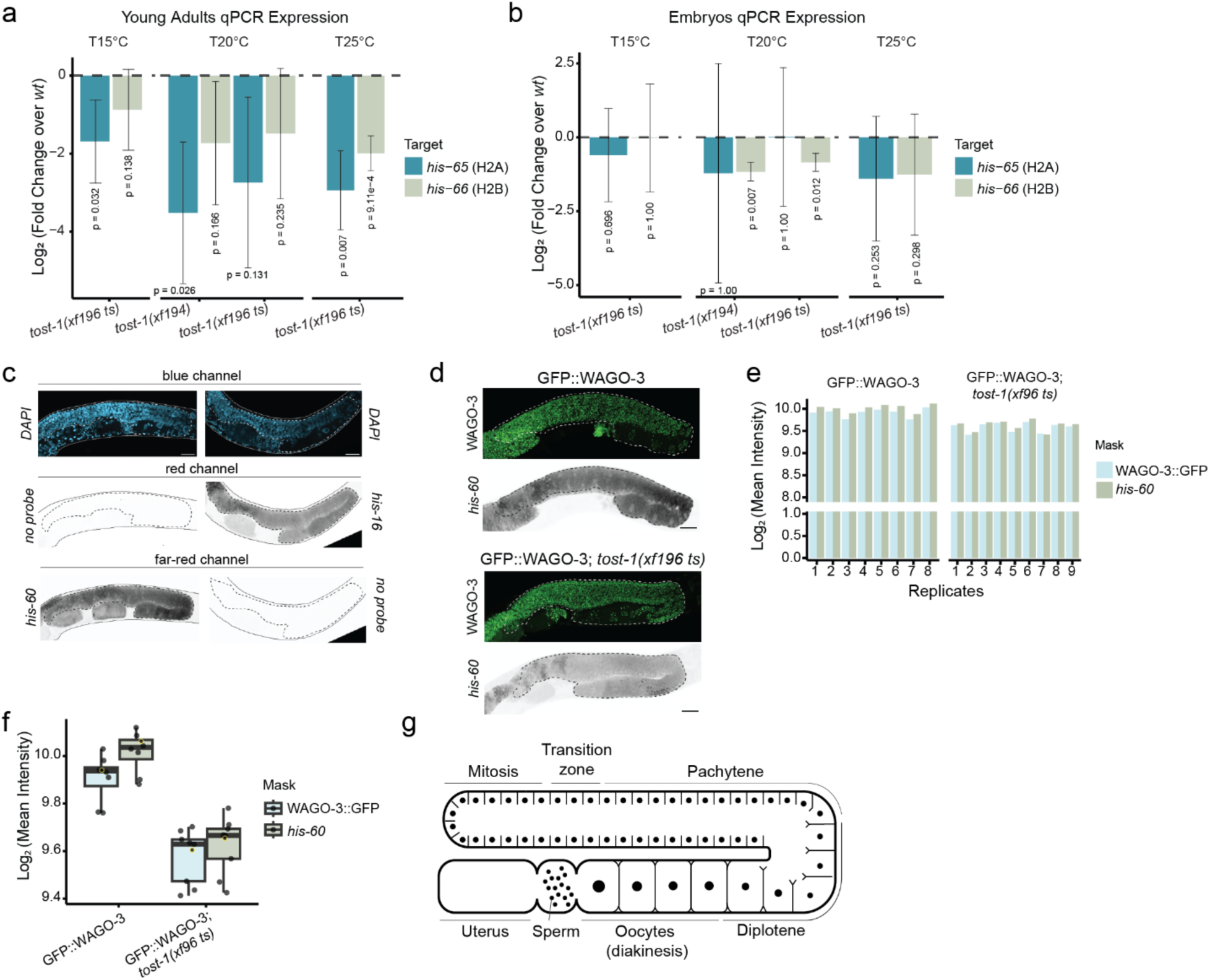
Histone mRNA levels are reduced in *tost-1* mutants. **a,** RT-qPCR analysis of *his-65* and *his-66* expression in young adults. Log2 fold changes relative to *wild-type* shown for *tost-1(xf196 ts)* at 15°C (n=3), 20°C (n=2), and 25°C (n=3), and *tost-1(xf194)* at 20°C (n=3). *Wild-type* controls: 15°C (n=3), 20°C (n=3), 25°C (n=3). Error bars represent 95% confidence intervals of log2 fold changes. P values calculated using Welch’s t-test with Bonferroni correction for multiple testing within each temperature. **b,** RT-qPCR analysis of *his-65* and *his-66* expression in embryos. Sample sizes for *tost-1(xf196)*: 15°C (n=3), 20°C (n=3), 25°C (n=3); *tost-1(xf194)*: 20°C (n=2). Wild-type controls: 15°C (n=3), 20°C (n=3), 25°C (n=3). Analysis as in panel a. **c,** smFISH specificity control showing *his-16* and *his-60* probe inverted fluorescence signals detected in red and far-red channels, respectively. **d,** smFISH of young adult worms expressing GFP::WAGO-3 in *wild-type* (n=8) or *tost-1(xf196 ts)* mutant background (n=9), showing GFP::WAGO-3 and *his-60* mRNA expression (inverted fluorescence signals). Scale bars, 10μm. **e,** Log2 mean intensity measurements of *his-60* smFISH signal in individual replicates from both strains. Region of interest for gonad measurements defined using either GFP::WAGO-3 signal or *his-60* smFISH signal as mask. **f,** Box plots of log2 mean intensity data from panel e. Yellow data points indicate samples corresponding to representative images shown. **g,** illustration of *C. elegans* adult gonad.

**Extended Data Figure 4.**
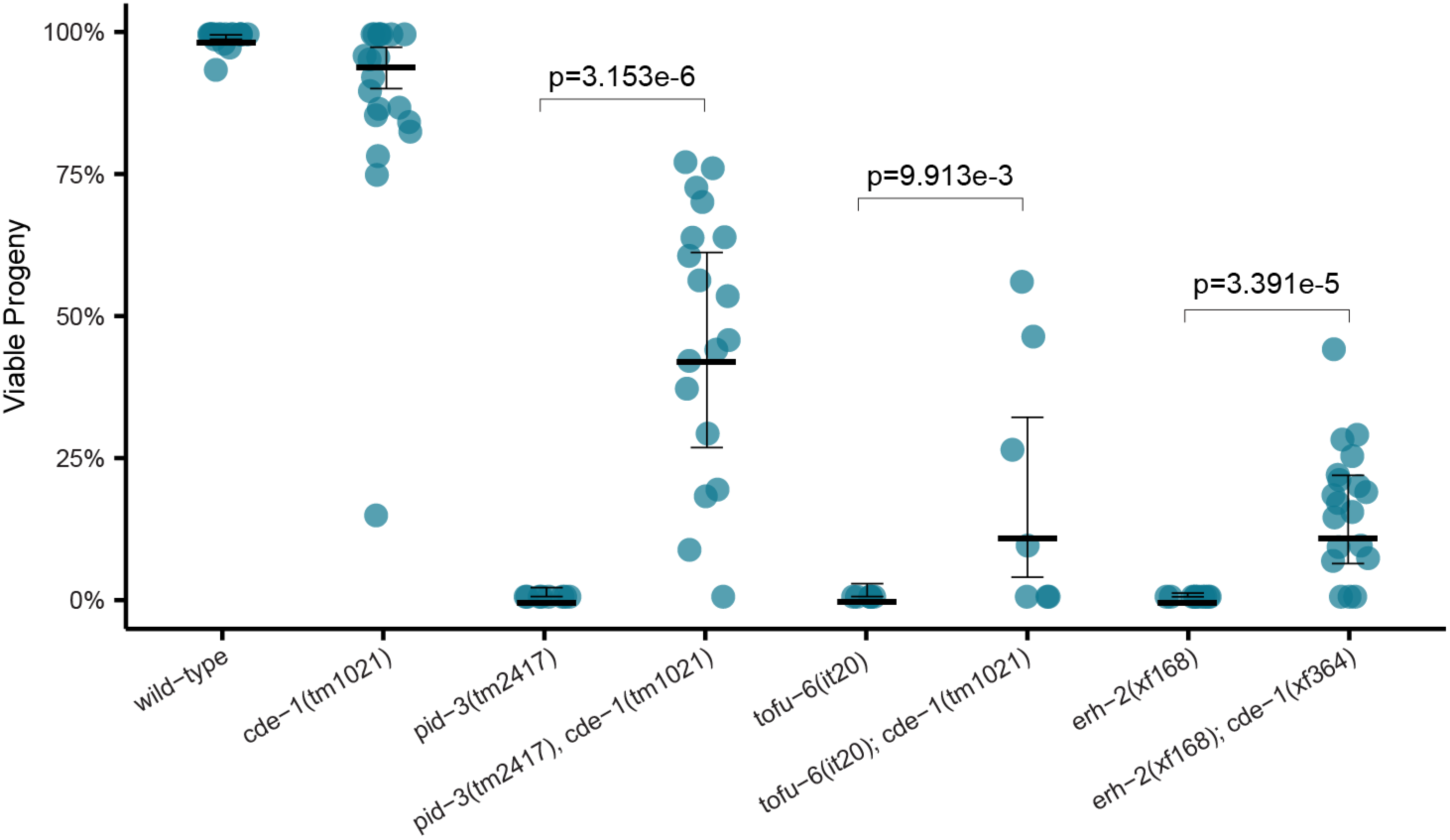
Loss of CDE-1 rescues embryonic lethality across multiple PETISCO components. Embryonic viability at 20°C for *wild-type* (n=18), *cde-1(tm1021)* (n=20), *pid-3(tm2417)* (n=8), *pid-3(tm2417); cde-1(tm1021)* (n=18), *tofu-6(it20)* (n=6), *tofu-6(it20); cde-1(tm1021)* (n=7), *erh-2(xf168)* (n=10), and *erh-2(xf168); cde-1(xf364)* (n=19). Statistical analysis using maximum a posteriori estimation within a Bayesian Generalised Linear Mixed-Effects Model with specific pairwise comparisons and multivariate-t method for multiple comparisons. Error bars represent 95% confidence intervals of the estimated viability.

**Extended Data Figure 5.**
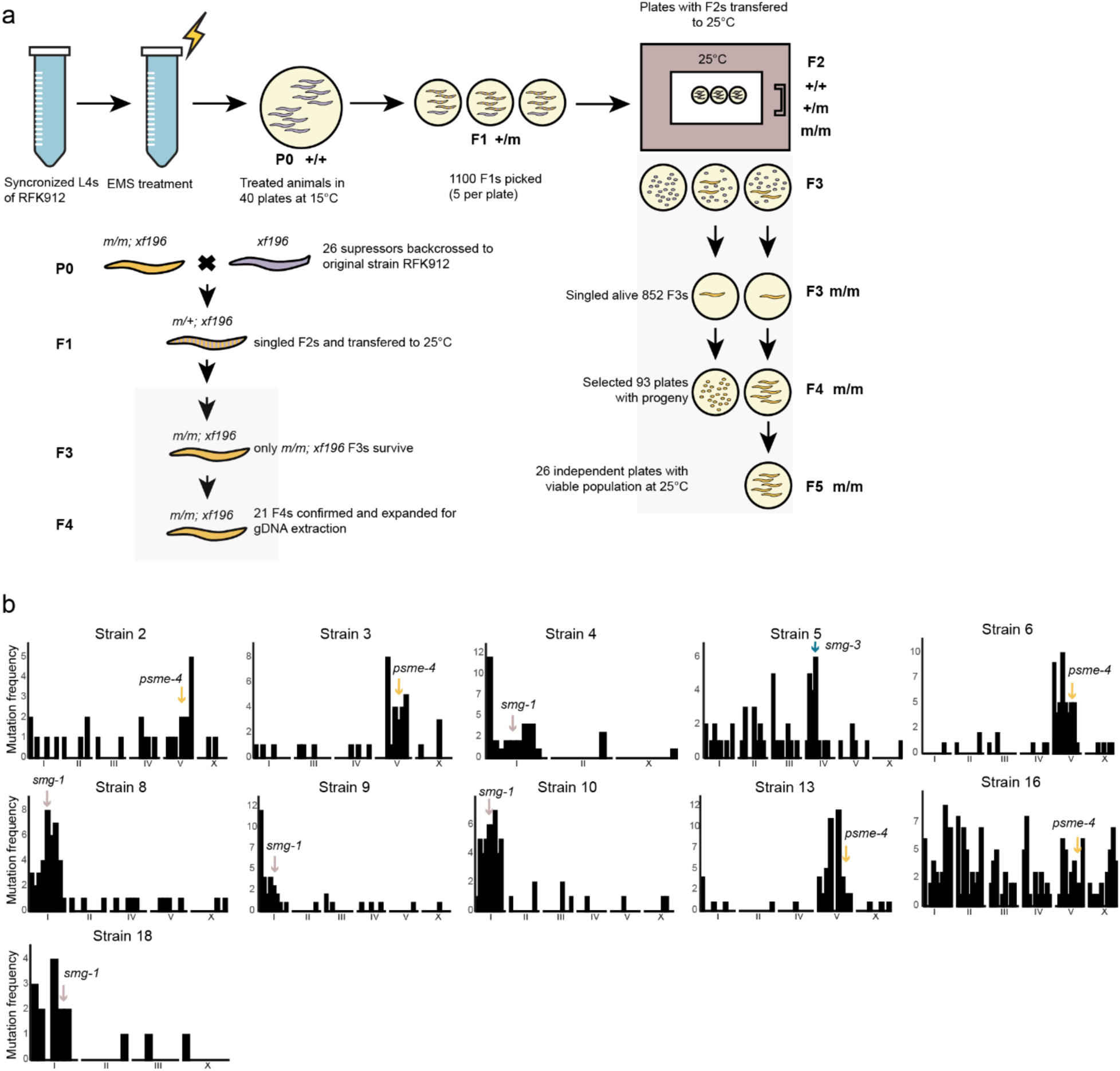
Forward genetic screen for suppressors of *tost-1* maternal-effect lethality. **a,** Schematic of EMS mutagenesis screen. Synchronised L4 RFK912 *tost-1(xf196 ts)* mutants’ larvae were treated with 47 mM EMS at 20°C for four hours, then plated onto 40 NGM plates and incubated at 15°C. After five days, 1100 F1 L4 larvae were picked (five per plate) and transferred to 25°C. Plates were screened after five days for F3 survivors (homozygous for suppressing mutations, m/m). A total of 852 F3 individuals were singled and maintained at 25°C for two generations, selecting viable progeny each generation. 26 suppressor strains with viable progeny at 25°C were identified and backcrossed with original RFK912 strain. After confirmation of survival at 25°C, 21 strains were expanded and subjected to whole-genome sequencing. **b,** Expected SNP (G/C-to-A/T) distribution across chromosomes in the strains with identified suppressors, indicated with an arrow. Chromosomes were divided into 10 bins.

**Extended Data Figure 6.**
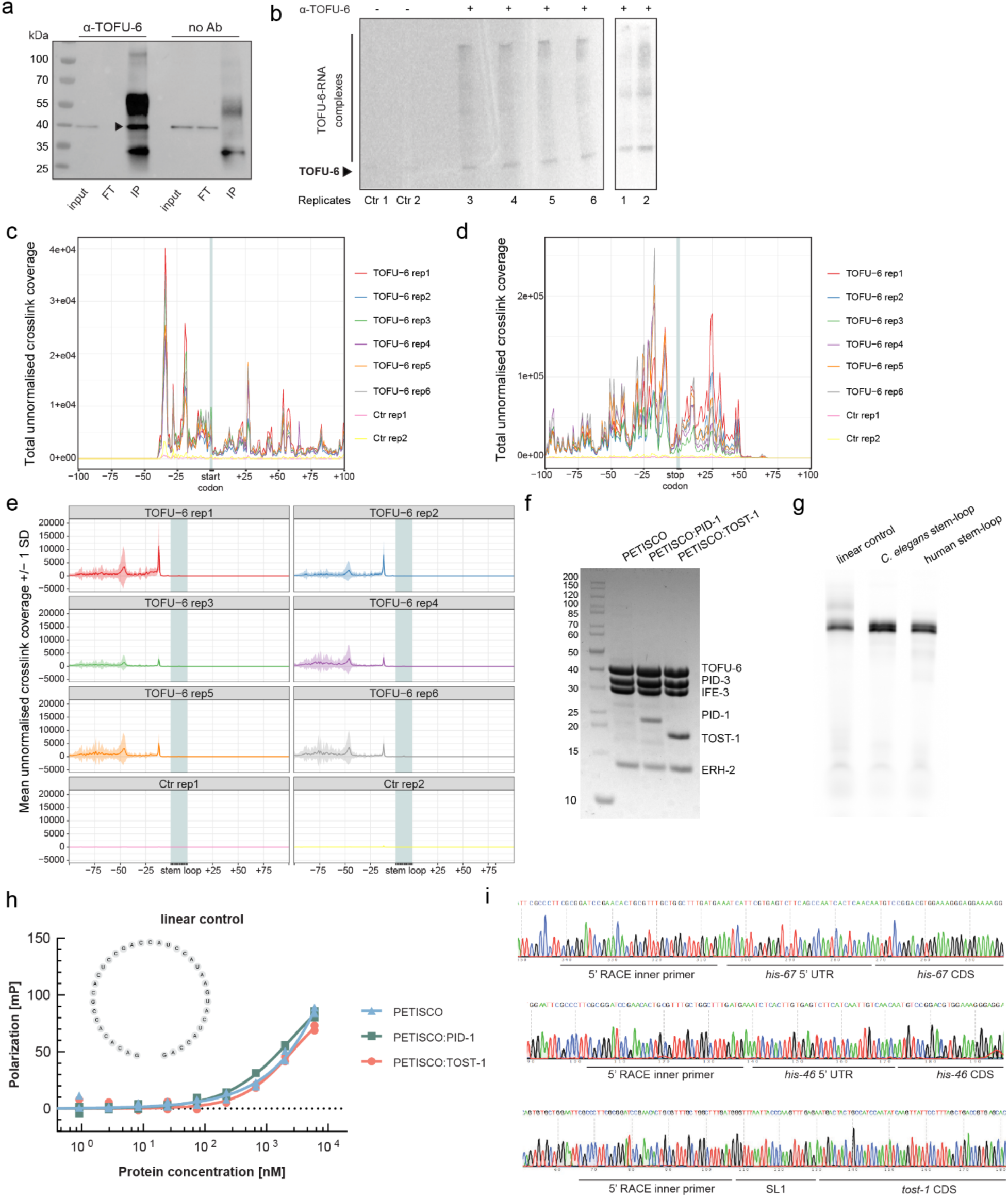
Direct binding of PETISCO:TOST-1 to histone mRNA stem-loop structures. **a,** Immunoprecipitation experiment using embryo lysates incubated with beads conjugated to anti-TOFU-6 antibody (α-TOFU-6) or unconjugated beads (no Ab). Input, immunoprecipitation (IP), and flow-through (FT) fractions analysed by western blot. Arrow indicates predicted TOFU-6 protein. **b,** Autoradiographs of final TOFU-6-RNA complexes used for library preparation (replicates 1-6). Samples prepared with unconjugated beads used as negative controls (Ctr rep1 and Ctr rep2). **c,d,** Meta-coverage plots showing the total coverage of crosslink sites relative to start codon (**c**) and stop codon (**d**) regions across RD histone genes in TOFU-6 iCLIP replicates. **e,** Meta-coverage plot visualising the mean coverage around the stem-loop region of RD histone genes for each replicate separately, shown with band of ±1 standard deviation (SD) around the mean. **c-e,** Data are shown unnormalised because the control libraries contained very low amounts of material, leading to low read numbers. Normalisation would artificially inflate control profiles to inappropriate levels. **f,** Coomassie-stained SDS polyacrylamide gel showing recombinant proteins: TOFU-6 (41.5 kDa), PID-3 (35.2 kDa), IFE-3 (28.2 kDa), PID-1 (19.4 kDa), TOST-1 (16.6 kDa), ERH-2 (13.3 kDa). **g,** Urea polyacrylamide gel showing labelled RNAs used for fluorescence polarisation experiments. RNAs are visualised by fluorescence. **h,** Fluorescence polarisation assays measuring binding of PETISCO alone, PETISCO:PID-1, and PETISCO:TOST-1 to fluorescently labelled linear RNA (n=2 replicates each). Structure of RNA substrate shown. **i,** Sanger sequencing results of representative clones from *his-67* gene (top), *his-46* gene (middle), and *tost-1* gene (bottom). Each sequence shows the position of the 5’ RACE inner primer located upstream of the 5’ sequence, which includes either the endogenous 5’ UTR or the splice leader SL1 sequence, followed by the start of the coding sequence (CDS) of the cloned cDNA.

